# DNA motif analysis of shear stress responsive endothelial enhancers reveals differential association of KLF and ETV/ETS binding sites with gained and lost enhancers

**DOI:** 10.1101/2021.09.21.460846

**Authors:** Roman Tsaryk, Nora Yucel, Elvin V. Leonard, Noelia Diaz, Olga Bondareva, Maria Odenthal-Schnittler, Zoltan Arany, Juan M. Vaquerizas, Hans Schnittler, Arndt F. Siekmann

## Abstract

Endothelial cells (EC) lining blood vessels are exposed to mechanical forces, such as shear stress exerted by the flowing blood. These forces control many aspects of EC biology, including vascular tone, cell migration and proliferation in addition to cell size and shape. Despite a good understanding of the genes and signaling pathways responding to shear stress, our insights into the transcriptional regulation of these responses is much more limited. In particular, we do not know the different sets of regulatory elements (enhancers) that might control increases or decreases in gene expression. Here, we set out to study changes in the chromatin landscape of human umbilical vein endothelial cells (HUVEC) exposed to laminar shear stress. To do so, we performed ChIP-Seq for H3K27 acetylation, indicative of active enhancer elements and ATAC-Seq to mark regions of open chromatin in addition to RNA-Seq on HUVEC exposed to 6 hours of laminar shear stress. Our results show a correlation of gained and lost enhancers with up- and downregulated genes, respectively. DNA motif analysis revealed an over-representation of KLF transcription factor (TF) binding sites in gained enhancers, while lost enhancers contained more ETV/ETS motifs. We validated a subset of flow responsive enhancers using luciferase-based reporter constructs and CRISPR-Cas9 mediated genome editing. Lastly, we characterized shear stress responsive genes in ECs of zebrafish embryos using RNA-Seq. Together, our results reveal the presence of shear stress responsive DNA regulatory elements and lay the groundwork for the future exploration of these elements and the TFs binding to them in controlling EC biology.

## Introduction

The formation of properly patterned blood vessels needs to be tightly controlled during embryonic development as they provide growing tissues with signaling molecules and the means of gas and metabolite exchange. Early vascular development is often stereotyped and depends on hard-wired genetic programs, while at later stages blood flow patterns provide additional mechanical cues^1^. These include tangential fluid shear stress and circumferential cyclic strain^2^. During angiogenesis stages, the formation of new sprouts is initiated in areas with lower shear stress^3^, suggesting that high levels of shear stress tone down angiogenesis and induce endothelial quiescence^4, 5^. Mechanical forces are also instrumental in vascular plexus remodeling that occurs through the pruning of excess blood vessels, ultimately resulting in the formation of mature and efficient vascular networks^6^. In this setting, shear stress induces the migration of ECs from vessels with low blood flow into vessels with high flow, as observed in the mouse retina^7^ and yolk sac vasculature^8^, zebrafish brain^9^, eye^10^ and subintestinal vessels^11^, as well as in quail embryos^12^.

Besides its role in vascular development and remodeling, shear stress is critical for blood vessel maintenance^13^. Unidirectional laminar shear stress affects proliferation and survival of ECs^14^ and has anti-thrombotic^15^ and anti-inflammatory effects^16^. It also controls endothelial barrier function^17^ and the cytoskeleton^18^. In contrast, oscillatory shear stress observed in areas of vessel curvature and vessel branching induces expression of pro-inflammatory endothelial markers^19^ and thus contributes to atherosclerosis development^20^. ECs can respond to changes in shear stress by regulating vessel size^21^. We have shown that in early embryonic development in zebrafish shear stress can induce changes in EC shape and orientation leading to vessel constriction^22^. In adult vessels, increases in blood flow can either result in vessel dilation through the release of nitric oxide (NO) from ECs^23^ or in vasoconstriction, mediated by the mechanosensing ion channel Piezo1^24^ The interplay of these two processes is likely important for blood pressure regulation.

Previous studies identified TFs of the Krüppel-like factor family as master regulators of the shear stress response^25, 26^, in particular KLF2^27^ and KLF4^28^. Fledderus et al. suggested that KLF2 together with nuclear factor erythroid-2 (Nrf2) regulates the expression of 70% of flow responsive genes^29^. However, other studies showed a lower magnitude of KLF2-dependent regulation^30^ and the association of other members of the KLF family or Sp1 to KLF binding sites^31^. Single cell ATAC-Seq identified binding sites for KLF4 and Nrf1 in ECs exposed to stable flow in carotid arteries, while ECs in arteries with disturbed flow showed enrichment of TEAD1 and ETV3 TF binding sites, among others^32^. We previously identified shear stress responsive elements in ECs exposed to atheroprone oscillatory shear stress and showed that activation of YAP/TAZ signaling was involved in mediating the EC response to this type of shear stress^33^. Work investigating histone modifications in response to pulsatile shear stress identified KLF4 as a downstream TF^34^. Furthermore, early growth response (EGR1) and SMAD1/5 have been implicated in the regulation of shear stress-induced gene expression changes^35, 36^, while the Snail TF responds to low shear stress in mediating endothelial to mesenchymal transition^37^. Therefore, our understanding of the transcriptional regulation of the shear stress response is far from complete and might involve the use of distinct sets of TFs depending on the flow regime and/or EC type examined^5^.

In this study, we used chromatin immunoprecipitation sequencing (ChIP-Seq) for acetylated H3K27, which marks active enhancers^38^, in combination with assay for transposase-accessible chromatin using sequencing (ATAC-Seq) to identify and characterize laminar shear stress responsive enhancer elements in cultured ECs. Our results show that proximity to gained or lost enhancers can faithfully predict shear stress induced changes in gene expression. We further identified binding sites for members of the KLF family of TFs in gained enhancers, while ETS binding sites were overrepresented in lost enhancers. We also identified several novel DNA motifs. Finally, deleting shear stress responsive enhancer elements using CRISPR-Cas9 mediated genome editing resulted in a reduction in shear stress regulation of SEMA7A and CXCR4 expression. Together, our results identified a suite of shear stress responsive enhancer elements and will thus serve as a resource for future studies investigating shear stress-mediated regulation of gene expression.

## Materials and Methods

### Cell Culture

To collect samples for ChIP-Seq, ATAC-Seq and RNA-Seq, HUVEC were isolated and cultured as described before^39^. The isolation was approved by the ethical board of the Westfaelische Wilhelms-Universitaet of Muenster and was in accordance with the Declaration of Helsinki. Within this framework, informed written consent was obtained from human subjects prior to HUVEC isolation. HUVEC were exposed to fluid shear stress of 18 dyn/cm^2^ in a BioTechFlow system (BTF system; MOS Technologies) at 37°C and under constant CO2 perfusion, as previously described^40^. HUVEC were seeded on 6-cm BTF glass plates coated with cross-linked gelatin and cultured to confluence (approximately 10^5^ cells/cm^2^) in EC growth medium (Promocell, Heidelberg, Germany). The maximum flow of 18 dynes/cm^2^ was achieved by computer-controlled cone rotation in culture medium containing 3% polyvinylpyrrolidone (PVP) within ten minutes (ramp function) and then kept constant for the required time, as described before^33^. Static controls were incubated in a cell culture incubator. For all other experiments, HUVEC were purchased (Gibco, # C0035C) and cultured in EBM-2 medium (Lonza, # CC3156) with EGM-2 supplements (Lonza, # CC-4176). HUVEC in passage 4 were seeded on ibiTreat 0.4 mm μ-slide I Lure (ibidi, # 80176) pre-coated with fibronectin (5 μg/ml; Sigma, # F2006), and exposed to shear stress (18 dyn/cm^2^) using an ibidi Pump System (ibidi, # 10902).

### Zebrafish Experiments

All zebrafish experiments were conducted on animals prior to the third day of embryonic development and are therefore not considered animal experiments. The maintenance of adult zebrafish for breeding purposes was in accordance with local and international guidelines and approved by the Landesamt fuer Natur, Umwelt und Verbraucherschutz Nordrhein-Westfalen (LANUV)^41^. Zebrafish were maintained as described previously^42^. *Tg(kdrl:H2B-EGFP)^mu122^* embryos^10^ were dechorionated at 48 hours post fertilization (hpf) using pronase and treated for 4 h with nifedipine (12.5 nM, Biomol, # Cay11106) and tricaine (1.3 mM, Sigma-Aldrich, # A5040) to stop the heart beat or with a corresponding amount of DMSO as a control. Embryos were used for RNA-Seq or in situ hybridization. For RNA-Seq 300 embryos per condition were collected in ice cold 10 % DMSO and kept at −80°C for 2 min. Afterwards, embryos were washed twice with 1x Hanks’ Balanced Salt Solution (HBSS, Gibco, # 14185-052) and deyolked with calcium-free Ringer’s solution. Embryos were then dissociated with trypsin-EDTA (0.5 %) and collagenase type IV (50 mg/ml). The reaction was stopped by addition of FCS (5 %) and cells were collected by centrifugation at 350 g for 5 min at 4°C and washed twice with 1x HBSS followed by centrifugation. After a final wash, cells were resuspended in 1x HBSS, passed through a 40 μm nylon filter to obtain a single cell suspension, which was subjected to fluorescence-activated cell sorting (FACS). EGFP positive ECs were collected in RLT buffer (RNeasy Micro kit, Qiagen).

### RNA-Seq

HUVEC or FACS-sorted zebrafish ECs were lysed with RLT buffer containing β-mercaptoethanol and RNA was isolated with the RNeasy Plus Mini or Micro Kit (Qiagen), respectively. RNA was quantified with NanoDrop (Thermo Fisher Scientific) and quality assessment was performed with a Bioanalyzer (Agilent). Libraries were prepared using TruSeq Stranded Total RNA Sample Prep with RiboZero depletion (Illumina), quantified with Qubit (Thermo Fisher Scientific), analyzed for fragment distribution with a Bioanalyzer and sequenced with paired-end settings (2 x 75 cycles) on a MiSeq System (Illumina).

### ChIP-Seq

ChIP was performed as previously described^33^. Briefly, HUVEC were cross-linked with formaldehyde (final concentration 1 %) for 5 min. After neutralization with glycine (0.125 M), cells were resuspended in nuclei lysis buffer and sonicated using an AFA sonicator (Covaris). Immunoprecipitation was performed using a H3K27ac antibody (Abcam, # ab4729) pre-bound to M-280 sheep anti-rabbit Dynabeads (Thermo Fisher Scientific, # 11204D). DNA was purified with phenol/chloroform/isoamyl alcohol and precipitated with ethanol. DNA was quantified using Qubit and fragment distribution was controlled with a Bioanalyzer. Libraries were prepared using TruSeq ChIP Sample Preparation Kit (Illumina), quantified by Qubit and analyzed for fragment distribution with a Bioanalyzer and used for paired-end sequencing (2 x 75 cycles) on a MiSeq System (Illumina).

### ATAC-Seq

HUVEC were collected via trypsinization and slowly frozen in FCS containing 10 % DMSO in an isopropanol chamber at −80°C according to a published protocol^43^. After thawing, cells were resuspended in EBM-2 medium and 50,000 cells were used for nuclei isolation and ATAC reaction and library preparation using the OMNI-ATAC protocol^43, 44^. Libraries were quantified with a NanoDrop and fragment distribution was analyzed on a Bioanalyzer. Libraries were used for paired-end sequencing (2 x 35 cycles) on a NextSeq500 system (Illumina).

### Data analysis

Sequencing reads were aligned to hg19 (human) or danRer10 (zebrafish) genomes. Two replicates per condition were analyzed.

#### RNA-Seq

Data was mapped using TopHat2 v2.1.1.^45^ and differential gene expression analysis between flow-treated samples and static controls was performed using DESeq2^46^. Read counts were obtained with HTSeq^47^ Genes were considered as differentially expressed when the FDR-adjusted p-value was < 0.05. Gene Set Enrichment Analysis (GSEA) was done at http://www.webgestalt.org and overrepresentation analysis (ORA) was performed with g:Profiler (https://biit.cs.ut.ee/gprofiler/gost).

#### ChIP-Seq

Sequencing reads from ChIP and input samples were mapped with Bowtie 2 v2.3.5.^48^ and Samtools v1.9.^49^ was used to edit the alignments. Reads were downsampled to equivalent read depth. Peak calling was performed with Genrich (https://github.com/jsh58/Genrich) applying the following parameters: genrich -v -e Y,MT -E blacklist.bed -t Sample1ChIP.bam, Sample2ChIP.bam -c Sample1Input.bam, Sample2Input.bam -o Sample.narrowPeak -k Sample.bedGraphish. Differential peak analysis was performed with DiffBind^50^ and peaks with FDR < 0.05 were used for further analysis. To analyse correlation between our and ENCODE replicates, Deeptools v2.29.2^51^ multiBamSummary was used to compute read coverages for genomic regions that were directly used to calculate and visualize pairwise Pearson’s correlation values.

#### ATAC-Seq

Sequencing reads were mapped with Bowtie2, downsampled to equivalent read depth and peak calling was performed with Genrich in ATAC-Seq mode using the following parameters: genrich -j -v -y -r -e Y,MT -E blacklist.bed -t Sample1.bam, Sample2.bam -o Sample.narrowPeak -k Sample.bedGraphish. Differential peak analysis was performed with DiffBind^50^ and peaks with FDR < 0.05 were used for further analysis.

ChIP-Seq and ATAC-Seq tracks were produced with deepTools bamCoverage, normalized to sequencing depth and visualized with UCSC Genome Browser. The overlap between ChIP-Seq and ATAC-Seq peaks was analysed with bedtools closest function and peaks within 100 bp of each other were considered to overlap. Heatmaps of overlapping regions were produced with deepTools. Peak association with neighbouring genes was performed with GREAT^52^.

Annotations of the peaks to defined genomic regions was done with HOMER^53^ annotatePeaks.pl and DNA motif analysis was performed with findMotifsGenome.pl function using default region size and corresponding common peaks as background regions.

### Luciferase assay

Putative enhancers were amplified from genomic DNA with Phusion High-Fidelity DNA Polymerase (New England Biolabs, # M0530). Betaine (1 M final concentration, Sigma Aldrich, # B0300) was added to the PCR reactions. Primers contained overhangs for cloning (primer sequences and overhangs are listed in Suppl. Table 1). PCR products were cloned into pCR4Blunt-TOPO vector (Invitrogen, # 450031) and the sequence of the cloned DNA was confirmed by Sanger sequencing. Inserts were cut out with appropriate restriction enzymes (Suppl. Table 1) and ligated with pre-digested pGL4.23[*luc*2/minP] (Promega, # E8411) plasmid upstream of a minimal promoter and a firefly luciferase gene. HUVEC were co-transfected using Lipofectamin 2000 (Invitrogen, # 11668027) and PLUS reagent (Invitrogen, # 11514015) with a firefly plasmid and the pRL-CMV plasmid (Promega, # E2261) containing the Renilla luciferase gene for normalization. HUVEC were exposed to shear stress (18 dyn/cm^2^ for 24 h) 2 h after transfection. Afterwards, cells were lysed and activity of firefly and renilla luciferase was measured with a Dual-Luciferase Reporter Assay System (Promega, # E1910).

### Enhancer activation with dCas9-p300

gRNA sequences were designed with chopchop (https://chopchop.cbu.uib.no) to target putative enhancers. Corresponding DNA oligonucleotides with BbsI overhangs (sequences are listed in Suppl. Table 1) were annealed and ligated with pre-digested pSPgRNA plasmid (Addgene, # 47108). HEK293T17 cells (ATCC, # CRL11268) were co-transfected using Lipofectamin 2000 (Invitrogen, # 11668027) and PLUS reagent (Invitrogen, # 11514015) with combinations of pSPgRNA plasmids and pcDNA-dCas9-p300Core plasmid (Addgene, # 61357) to activate putative SEMA7A enhancers^54^ SEMA7A expression was measured by qPCR 4 days after transfection. Alternatively, cells were co-transfected with SEMA7A-Luc4 and pRL-CMV (Promega, # E2261) constructs and enhancer activity was measured with a luciferase assay.

### CRISPR-Cas9-mediated enhancer removal

Putative enhancers were removed using CRISPR pairs. gRNA sequences were designed with chopchop (https://chopchop.cbu.uib.no) and corresponding DNA oligonucleotides containing BsmBI overhangs were annealed and ligated with lentiCRISPR v2 plasmid (Addgene, # 52961), which contains both Cas9 and sgRNA sequences^55^. The sequences targeting putative enhancers are listed in Suppl. Table 1. HEK293T17 cells (ATCC, # CRL11268) were transfected with specific lentiCRISPR v2 plasmids, as well as pMD2.G (Addgene, # 12259) and psPAX2 (Addgene, # 12260) plasmids using Fugene 6 Transfection Reagent (Promega, #E2311). 72 h after transfection, the supernatant was collected and filtered through a 0.45 μm filter (Sarstedt, # 83.1826) and virus was collected with LentiX Concentrator (Takara, # 631232). HUVEC were infected with combinations of two viruses and used 96 h after infection. Enhancer removal was confirmed by PCR, using Phusion High-Fidelity DNA Polymerase (New England Biolabs, # M0530), 1M betaine (Sigma Aldrich, # B0300) and the primers listed in Suppl. Table 1. PCR products were visualized on ethidium bromide agarose gels.

### qPCR

RNA was isolated with an RNeasy Micro Kit (Qiagen) and reverse transcribed using an iScript cDNA Synthesis Kit (Bio-Rad, # 1708891). Power SYBR Green PCR Master Mix (Applied Biosystems, # 4367659) was used to run qPCR in a 7900HT Fast Real-Time PCR System (Applied Biosystems). ddCt method was used to calculate relative expression using RPL13A or TBP as endogenous controls. Statistical analysis was performed on dCt data. Primer sequences are listed in Suppl. Table 1.

### ChIP-qPCR

DNA isolated after H3K27ac ChIP was used for qPCR with Power SYBR Green PCR Master Mix and primers listed in Suppl. Table 1. In addition, qPCR was performed on DNA from input control and H3K27ac enrichment was calculated as % of input.

### siRNA

HUVEC were transfected with 50 nM of siRNA using Oligofectamine (Invitrogen, # 12252011) and used for flow experiments in an ibidi system 48 h after transfection. Silencer Select Negative Control No. 1 siRNA (Invitrogen, # 4390843) and Silencer Select KLF2 siRNA (Invitrogen, # 4392420, ID # s20270) were used.

### Statistical analysis

Statistical analysis was performed with GraphPad Prism.

## Results

### Shear stress induces gene expression changes in ECs

To understand the regulation of gene expression by shear stress at the level of cis-regulatory elements we first performed RNA-Sequencing (RNA-Seq) in HUVEC exposed to shear stress in a BioTechFlow system (Fig. 1A) and compared gene expression to cells cultured under static conditions. To determine the optimal timepoint for our analysis, we initially exposed HUVEC to 18 dyn/cm^2^ for 30 min and for 6 h. After 30 mins of shear stress, 59 genes were upregulated and 17 were downregulated (Suppl. Fig. 1, Suppl. Data 1). Several chaperones and co-chaperones (HSPA1A, HSPA1B, HSP6, DNAJB1) were the most highly upregulated genes. In addition, the known shear stress-regulated TFs KLF2 and KLF4 were upregulated, pointing to the onset of a shear stress-specific transcriptional program already after a short exposure. After 6 h of shear stress, 1014 genes were differentially expressed (DE) with 647 genes up-regulated and 367 down-regulated when compared to control (Fig. 1B, Suppl. Data 1). We detected differential expression of known flow-responsive genes, such as KLF2, KLF4, NOS3, CYP1B1, WNT9B, which were upregulated by shear stress, and EGR1 and CXCR4, which were downregulated by shear stress (Fig. 1B, Suppl. Fig. 2A, B). Based on the results of these initial experiments and a separate longitudinal study^56^, we decided to continue our analysis with 6 h of shear stress exposure. We first validated our RNA-Seq results using several methods. Quantitative polymerase chain reaction (qPCR) on the most up- and downregulated DE genes and of nonregulated genes with a Log2 fold change close to 0 revealed a high degree of correlation between RNA-Seq and qPCR results (Suppl. Fig. 2C). Pathway analysis of DE genes using gene set enrichment analysis (GSEA, Fig. 1C) and overrepresentation analysis (ORA, Suppl. Fig. 2D) revealed that “fluid shear stress and atherosclerosis” was the most enriched pathway in our dataset. We also compared our RNA-Seq dataset with published studies on shear stress-induced gene expression changes in ECs (Suppl. Fig. 2E). This analysis showed a 34.5 % overlap with all DE genes from an RNA-Seq dataset of HUVEC exposed to 20 dyn/cm^2^ for 3 days by Maleszewska et al.^57^ and a 32.5 % overlap with all DE genes from a meta-analysis of multiple microarray-based reports study by Maimari et al.^58^. Additionally, we detected a substantial overlap (15.7 %) between our data and the dataset generated by Dekker et al.^4^, mimicking the shear stress response through KLF2 overexpression (Suppl. Fig. 2E). Together, our RNA-Seq data are highly reproducible and partially match previously published datasets, while at the same time revealing considerable differences in the transcriptional response of ECs to different flow regimes. They furthermore suggest that TFs other than KLF2 play important roles in regulating EC flow responses.

**Figure 1.**
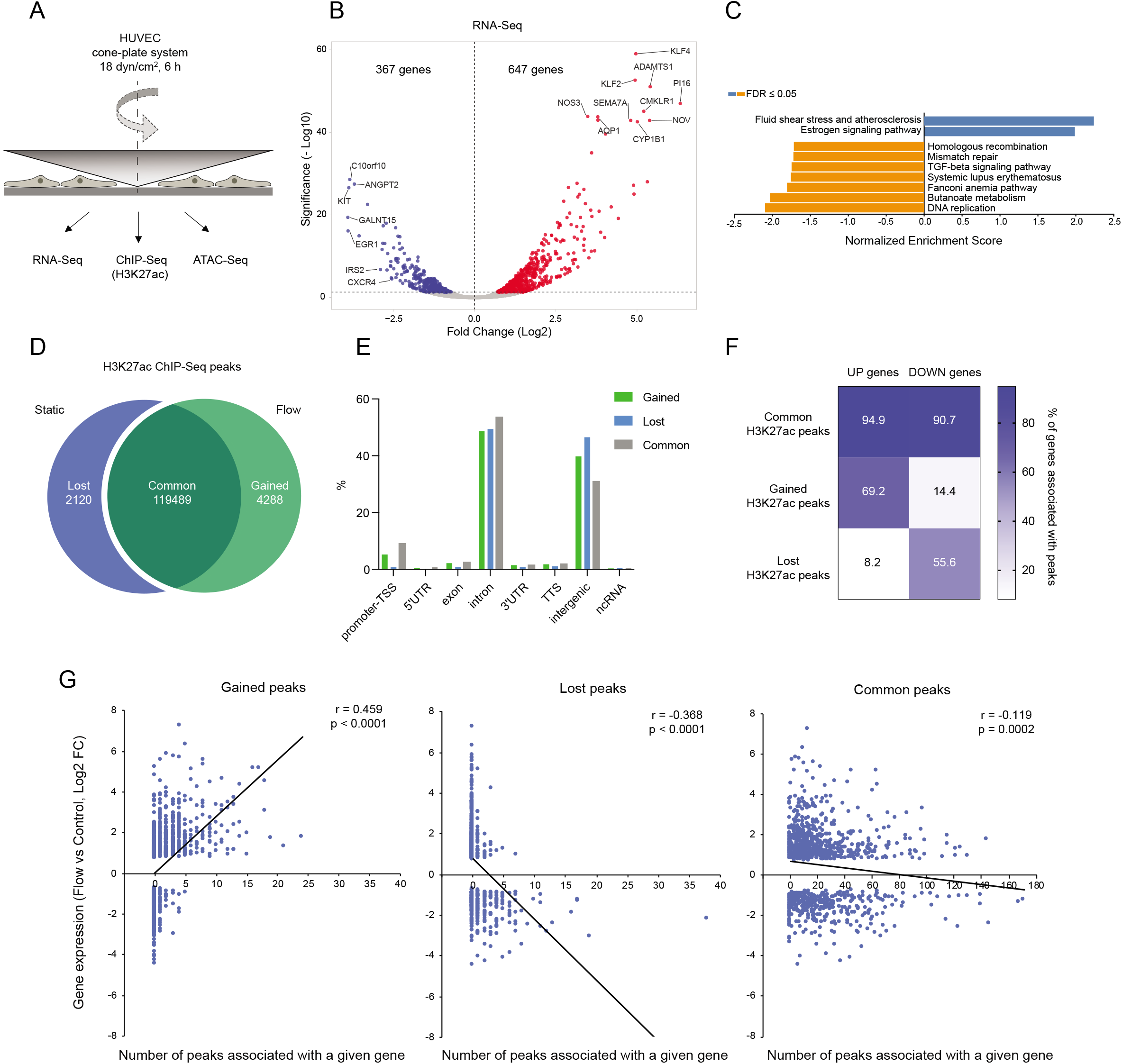
Shear stress regulates the expression of a defined set of genes that correlates with enhancer activation. **A**. HUVEC exposed to unidirectional shear stress (18 dyn/cm^2^) for 6 h were used for RNA-Seq, ATAC-Seq and H3K27ac ChIP-Seq. **B**. Volcano plot showing gene expression changes in HUVEC under shear stress. Differentially expressed (DE) genes (FDR < 0.05) are labelled in red (upregulated) and blue (downregulated under shear stress). **C**. Gene Set Enrichment Analysis (GSEA) of KEGG pathways regulated in the RNA-Seq dataset. **D**. The number of H3K27ac ChIP-Seq peaks in HUVEC under static conditions (“lost” after shear stress) or exposed to shear stress (“gained” after shear stress) and the peaks “common” for both conditions. **E**. Annotation of gained, lost and common H3K27ac peaks to defined regions of the genome. **F**. Heatmap representing the spatial association of the genes up- or down-regulated by shear stress with identified common, gained or lost H3K27ac ChIP-Seq peaks. Numbers represent the percentage of up-or down-regulated genes, associated with at least one peak. **G**. Comparison of gene expression (log2 fold change (FC) from RNA-Seq) to the number of gained, lost or common H3K27ac peaks spatially associated with a particular gene. Pearson correlation coefficient (r) and p-value of the correlation are shown.

### ChIP-Seq identification of flow responsive regulatory elements marked by H3K27ac

To interrogate which enhancer elements might be responsible for the observed changes in gene expression, we performed ChIP-Seq analysis of shear stress (18 dyn/cm^2^ for 6 h) exposed and unexposed static control HUVEC using an antibody specific for histone 3 acetylated at the lysine 27 residue (H3K27ac), which marks active enhancers^38^. To validate our data, we compared our data with previously generated datasets from the ENCODE consortium^59^. Our analysis revealed a high degree of correlation between our replicates, both for static and shear-stress-exposed conditions (Suppl. Fig. 3A). Our static replicates correlated well with HUVEC samples from ENCODE (Pearson coefficient 0.72 – 0.82), similar to the correlation between individual ENCODE replicates (0.78 – 0.89). The analysis of our ChIP-Seq data identified 119489 peaks (DNA regions with significantly enriched H3K27ac signal compared to input control) common for both flow and static conditions (Fig. 1D). Furthermore, we identified 4288 peaks specific for shear stress-exposed cells (“gained” peaks) and 2120 peaks present only in control cells (peaks “lost” in shear stress condition; Fig. 1D). Annotation of gained and lost H3K27ac peaks to defined regions of the genome revealed that most of the peaks were in introns of coding genes and in intergenic regions, while a relatively low number of peaks was associated with gene promoters (Fig. 1E). To further validate our results, we performed qPCR with primers for selected regions showing a high enrichment for H3K27ac. These data on independent biological replicates confirmed the results from our ChIP-Seq dataset (Suppl Fig. 3B). Thus, we obtained a validated dataset of flow-responsive gene regulatory elements.

### Flow responsive cis-regulatory elements associate with flow regulated genes

As proximity favors enhancer-promoter choice to some extent^60, 61^, we associated H3K27ac peaks to neighboring genes using the GREAT algorithm^52^. This algorithm identifies the nearest genes of a given peak within a range of 1 Mb. We then compared these genes to the set of DE genes from our RNA-Seq data. This analysis revealed that upregulated genes were more often associated with gained peaks, while downregulated genes were more often associated with lost peaks. Hence, 69 % of upregulated genes associated with at least one gained peak, while this number was only 14 % for downregulated genes. Conversely, 56 % of downregulated genes associated with at least one lost peak, while only 8 % of upregulated genes had associated lost peaks (Fig. 1F). Upregulated genes had more gained peaks per gene associated with them, while downregulated genes associated with more lost peaks per gene. Additionally, the number of associated peaks correlated with gene expression levels (Fig. 1G). This correlation was not observed for common H3K27ac peaks (Fig. 1F, G). These data are in line with our hypothesis that shear stress activates enhancers associated with upregulated genes and inhibits enhancers associated with downregulated genes. They furthermore show that our ChIP-Seq analysis has predictive power to anticipate shear stress-mediated regulation of gene expression.

### Analysis of chromatin accessibility associated with shear stress-responsive regulatory elements

To analyze chromatin accessibility, which is often associated with TF binding, we performed ATAC-Seq in HUVEC exposed to shear stress and compared the data to our sets of DE genes and H3K27ac peaks. This analysis identified over 104000 common ATAC peaks, while 7661 peaks were gained, and 1027 peaks were lost upon shear stress exposure of the cells (Fig. 2A). Like H3K27ac peaks, gained and lost ATAC-Seq peaks were preferentially located in gene introns and intergenic regions (Suppl. Fig. 4A). To understand how chromatin accessibility correlated with histone acetylation, we analyzed the overlap between ATAC-Seq and H3K27ac peaks (located within 100 bp of each other, Fig. 2B). 21 % of gained ATAC-Seq peaks were associated with gained H3K27ac peaks, while 24 % of lost ATAC-Seq peaks overlapped with lost H3K27ac peaks (Suppl. Fig. 4B), indicating a similar mode of activation/deactivation for this subset of peaks. The numbers of ATAC-Seq peaks associated with H3K27ac peaks showing an opposite mode of regulation was lower. Only 0.1 % of gained ATAC-Seq peaks overlapped with lost H3K27ac peaks and only 0.2 % of lost ATAC-Seq peaks overlapped with gained H3K27ac peaks. A substantial amount of both gained and lost ATAC-Seq peaks corresponded with common H3K27ac peaks or showed no overlap. A reverse analysis of the overlap between H3K27ac peaks and ATAC-Seq peaks showed a similar tendency. The overlap was higher for gained H3K27ac and ATAC-Seq peaks and for lost H3K27ac and ATAC-Seq peaks (45 % and 15 %, respectively) than vice versa (0.1 % for both, Suppl. Fig. 4C). This analysis also revealed that many H3K27ac peaks were associated with common ATAC-Seq peaks (48 % of the gained and 51 % of the lost H3K27ac peaks, Suppl. Fig. 4C). Therefore, our comparative analysis of ATAC-Seq and H3K27Ac peaks revealed the presence of distinct subsets of H3K27ac peaks. In the first set, changes in histone acetylation coincide with changes in chromatin accessibility, suggesting enhancer activation/deactivation in response to shear stress. A second, large subset of peaks contains regions of open chromatin, which do not change with changes in histone acetylation (Suppl. Fig. 4B). This could suggest the presence of poised enhancers that get activated by shear stress and active enhancers that get deactivated but remain poised. Lastly, there is a subset of H3K27ac peaks which do not overlap with ATAC-Seq peaks. It remains to be determined, whether these are functional enhancers or whether they are the result of limitations in our study.

**Figure 2.**
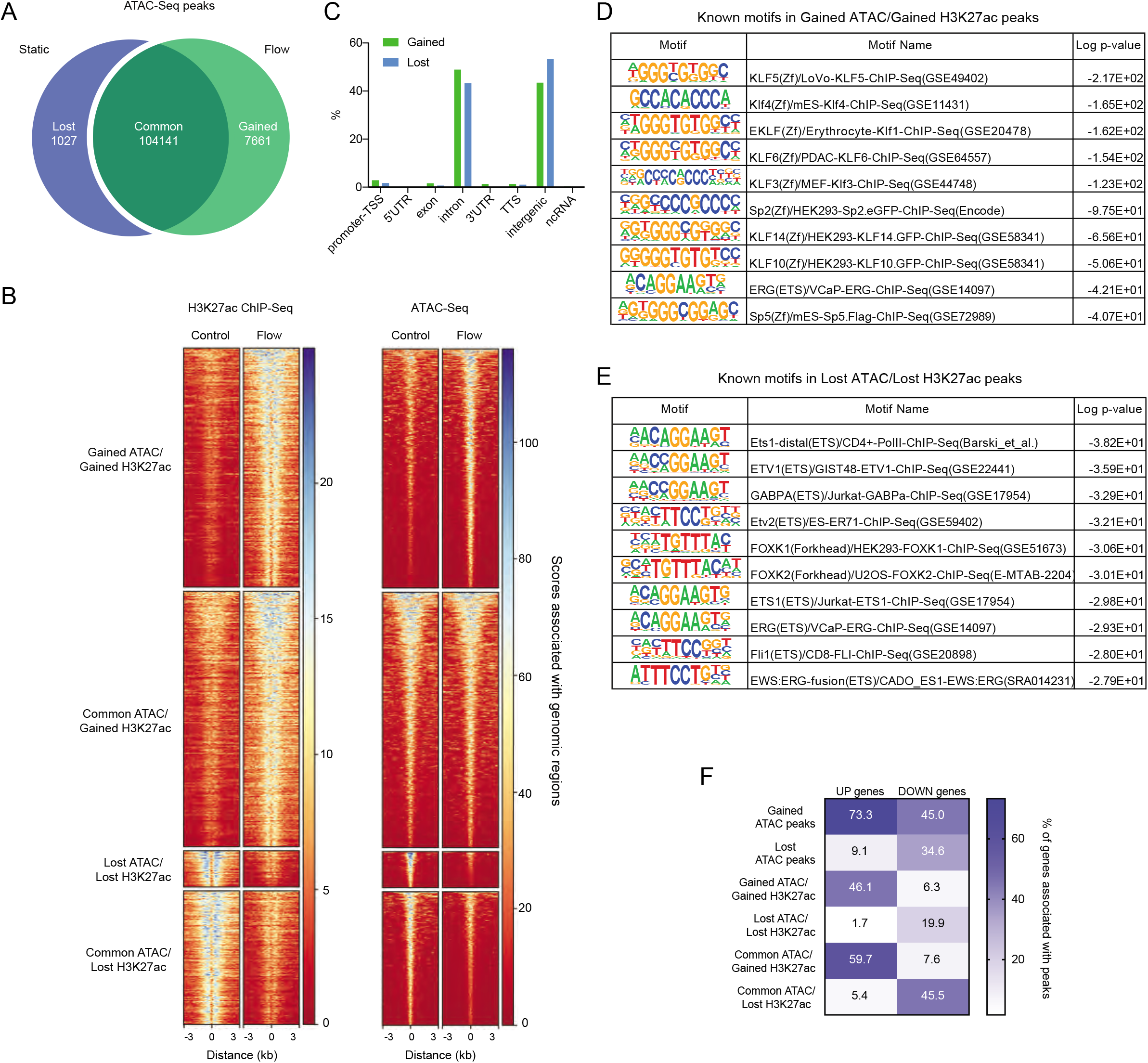
ATAC-Seq identifies DNA motifs that associate with shear stress responsive enhancers. **A.** Number of ATAC-Seq peaks specific for HUVEC under static conditions (“lost” after shear stress) or exposed to shear stress (“gained” after shear stress), as well as “common” for both conditions. **B**. Heatmap representing scores associated with genomic regions centered on the indicated H3K27ac- and ATAC-Seq peaks in control and flow (18 dyn/cm^2^ shear stress for 6 h) conditions. **C**. Annotation of gained and lost ATAC-Seq peaks that overlap with gained and lost H3K27ac ChIP-Seq peaks, respectively, to defined regions of the genome. **D, E**. Analysis of known DNA motifs enriched in gained (**D**) or lost (**E**) ATAC-Seq peaks that overlap with gained or lost H3K27ac ChIP-Seq peaks, respectively. The 10 most enriched motifs are shown. **F**. Heatmap representing spatial association of genes up- or down-regulated by shear stress with different subsets of ATAC-Seq peaks (all gained and lost ATAC-Seq peaks, gained and lost ATAC-Seq peaks that overlap with gained or lost H3K27ac ChIP-Seq peaks, respectively, or common ATAC-Seq peaks that overlap with gained or lost H3K27ac ChIP-Seq peaks). The numbers represent the percentage of up-or down-regulated genes, associated with at least one peak.

To identify TF binding site motifs in flow responsive enhancers, we analyzed the subset of H3K27ac peaks, which showed similar changes in H3K27ac and chromatin accessibility. These peaks were mostly located in introns and intergenic regions (Fig. 2C). In line with the role of KLF TFs in EC responses to shear stress, a number of KLF motifs were enriched in gained peaks (e.g. KLF5, KLF4, KLF1, KLF3, KLF6, KLF10, KLF14, Fig. 2D, Suppl. Fig. 4D; of note, a motif for KLF2 is not included in the database). Lost peaks were enriched in motifs for endothelial lineage TFs ETS1, ETV1, ETV2 and FLI1 (Fig. 2E, Suppl. Fig. 4E). Interestingly, common ATAC peaks that overlapped with gained H3K27ac peaks also contained several enriched KLF motifs (Suppl. Fig. 4F). However, other motifs such as JUNB, FOSL2 and ETV2 were also highly enriched in this subset. Motifs found in common ATAC peaks that overlapped with lost H3K27ac peaks were similar to those found in lost ATAC peaks overlapping with lost H3K27ac peaks (compare motifs in Suppl. Fig. 4G with those in Fig. 2E). Together, our analysis uncovered several known and novel DNA motifs enriched in shear stress activated enhancers.

### Changes in gene expression correlate with chromatin accessibility

To understand how chromatin accessibility predicts changes in gene expression, we associated ATAC-Seq peaks with nearby genes using the GREAT algorithm. Comparing genes associated with ATAC-Seq peaks with DE genes from our RNA-Seq dataset revealed that upregulated genes associated more with gained than lost ATAC-Seq peaks and vice versa (Fig. 2F). This association was more obvious for upregulated genes, and it was higher when considering all ATAC-Seq peaks compared to the ones that overlap with H3K27ac peaks. Interestingly, analysis of the genes associating with H3K27ac peaks that overlap with common ATAC peaks revealed that they also had a high degree of association with DE genes. Hence, 60% of upregulated genes had gained H3K27ac peaks that overlapped with common ATAC peaks in their vicinity and 46% of downregulated genes associated with lost H3K27ac peaks that overlapped with common ATAC-Seq peaks (Fig. 2F). Together, our data might indicate the existence of different modes of regulation of shear stress-induced gene expression.

### Functional analysis of identified putative enhancers

Previous studies have shown that only a fraction (26%) of bioinformatically identified enhancer elements are functionally relevant in reporter assays^62^, while even fewer showed activity in large scale functional screens using the KRAB repressor^61, 63^. Therefore, the flow responsiveness of identified putative enhancers and their regulation of flow-responsive genes need to be validated experimentally. We tested flow responsiveness of several putative enhancers using luciferase assays. To do so, we cloned putative enhancers upstream of a minimal promoter regulating the expression of firefly luciferase (Suppl. Fig. 5A). Out of 14 constructs tested, 3 showed a significant increase in luciferase activity in response to shear stress (18 dyn/cm^2^ for 24 h; Suppl. Fig. 5B). This correlated with higher H3K27ac in these regions and with their association with upregulated DE genes. Most DE genes associated with multiple peaks. Since their cooperation might be necessary for shear stress-induced gene expression, putative enhancers may not function in isolation in a luciferase assay.

We then concentrated on the SEMA7A gene, which is associated with several gained H3K27ac peaks (in the promoter region and 45-55 kb downstream of the transcription start site). These overlapped with both gained and common ATAC peaks (Fig. 3A). This correlated with SEMA7A expression, which was induced 35-fold in shear stress exposed HUVEC (Fig. 1B, Suppl. Fig. 2A), while qPCR showed a 41-fold induction (Fig. 3B). SEMA7A was also upregulated in several published datasets of flow-regulated genes^57, 58^. To confirm enhancer activity of putative SEMA7A enhancers, we cloned 4 luciferase constructs (Luc 1 - 4; Fig. 3A) and tested them for flow response. Construct Luc2, containing the sequence in the promoter region, showed stronger activity compared to an empty vector (data not shown), but no flow response (Fig. 3C). In contrast, Luc4 showed a strong activation in response to shear stress with relatively high basal activity observed also under static conditions confirming its flow-responsive enhancer activity (Fig. 3C). We confirmed H3K27ac enrichment in this flow-responsive enhancer by ChIP-qPCR (Fig. 3D). Luc1 and 3 showed neither basal, nor flow-induced activity. Since the activity of EC-specific enhancers has been reported to be higher in conjunction with their own promoters compared to minimal promoters^64^, we combined Luc2 and Luc4 in one construct (Luc2+4). Indeed, the flow response of Luc2+4 was higher than that of Luc4 alone (Suppl. Fig. 5E), suggesting cooperative interactions between this enhancer and the SEMA7A promoter. To further investigate this enhancer, we used dCas9-p300 to target and activate the Luc4 enhancer using sets of gRNAs (Suppl. Fig. 5C, see Fig. 3A for gRNA targeting location) specific for different regions of the enhancer in HEK293T cells. These cells are easier to transfect compared to ECs and they have lower SEMA7A expression. HEK293T cells were transfected with corresponding gRNA-coding plasmids together with dCas9-p300 and the Luc4 plasmid. We detected a 50-fold increase in luciferase activity when using 12 gRNAs targeting the whole length of the Luc4 sequence (4 kb) compared to cells transfected with Luc4 and control plasmids indicating successful targeting of the SEMA7A enhancer (Suppl. Fig. 5F). Targeting the endogenous enhancer with the same gRNAs resulted in 3.5-fold SEMA7A mRNA upregulation in HEK293T cells without shear stress (Fig. 3E), with different sets of gRNAs showing similar activities as in luciferase assays. Importantly, targeting another region downstream of SEMA7A with no H3K27ac ChIP-Seq and ATAC-Seq peaks led to a lower change in SEMA7A expression (gRNAs N1-4; Fig. 3A). In a complementary approach, we deactivated the SEMA7A enhancer using Cas9 and different pairs of gRNAs to remove parts of the enhancer (Suppl. Fig. 5D; Fig. 3F, G). Exposing targeted cells to shear stress resulted in a smaller induction in SEMA7A expression (Fig. 3H). We therefore identified a gained flow-responsive enhancer that can regulate SEMA7A expression.

**Figure 3.**
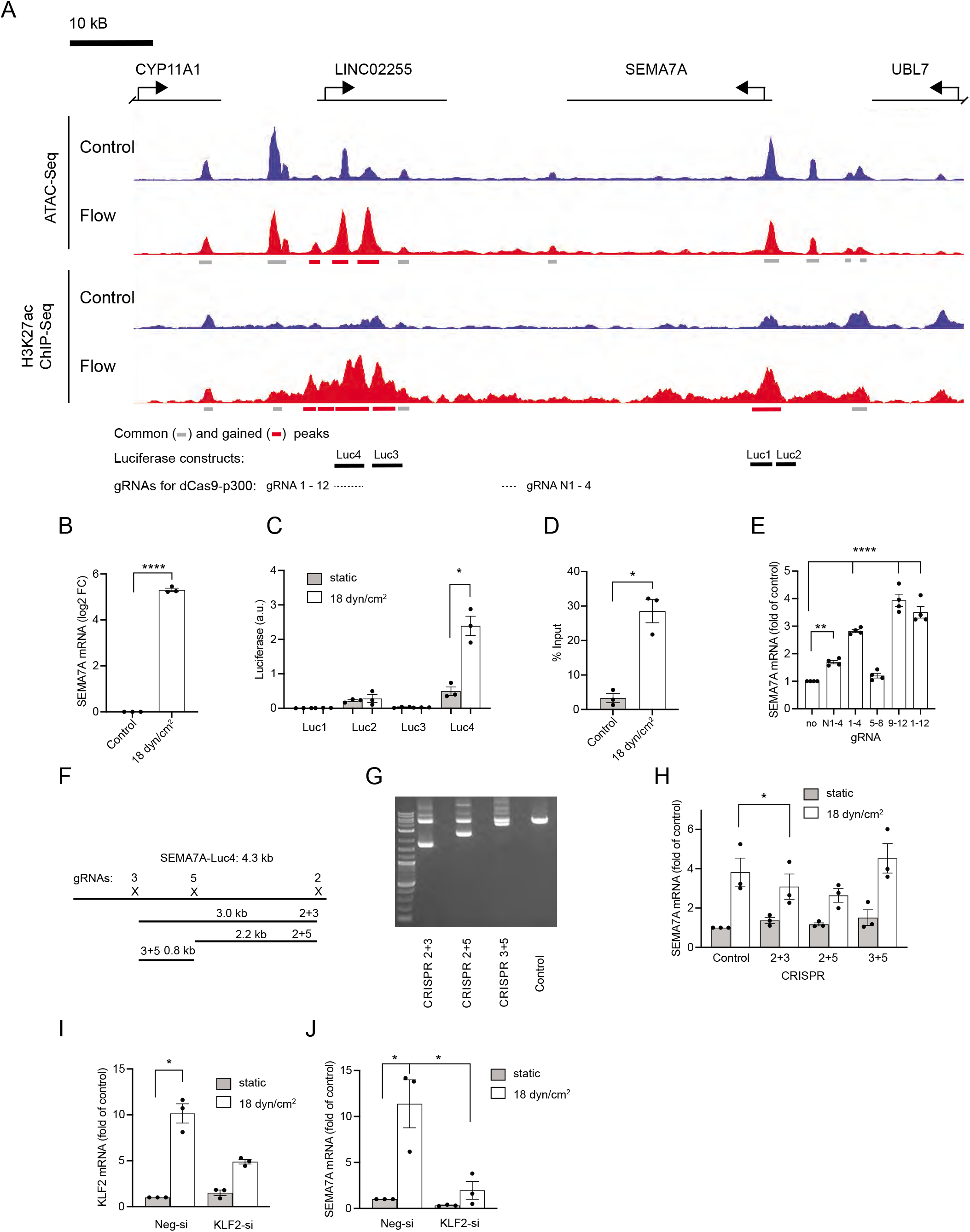
Identification and characterization of a shear stress responsive enhancer in the SEMA7A gene. **A.** Normalized ATAC-Seq and H3K27ac ChIP-Seq tracks in the vicinity of the SEMA7A gene. Bars below ATAC and ChIP-Seq tracks show identified common (grey) and gained (red) ATAC-Seq or H3K27ac ChIP-Seq peaks. In addition, the regions used for luciferase constructs in **C** and locations targeted by gRNAs in **E** are shown underneath the tracks. **B**. SEMA7A mRNA expression in HUVEC exposed to 18 dyn/cm^2^ for 6 h (Log2 fold of control, mean ± sem, n = 3, **** p < 0.001, two-tailed paired t-test). **C**. Luciferase assay showing luciferase activity of the constructs containing DNA regions indicated in **A** upstream of firefly luciferase in HUVEC exposed to shear stress (18 dyn/cm^2^ for 24 h, data are presented in arbitrary units (a.u.) after normalization to the signal of a renilla luciferase construct; mean ± sem, n = 3, * p < 0.05, two-tailed paired t-test). **D**. ChIP-qPCR showing higher H3K27ac ChIP-Seq enrichment in the region corresponding to Luc4 (see **A**) in HUVEC exposed to shear stress (18 dyn/cm^2^ for 6 h, data is presented as % input, mean ± sem, n = 3, * p < 0.05, two-tailed paired t-test). **E**. SEMA7A mRNA expression in HEK293T17 cells transfected with plasmids encoding dCas9-p300 and different gRNAs (combination of 4 or 12 gRNAs) targeting genomic regions downstream of the SEMA7A gene (marked in **A**; data is shown as fold of control, mean ± sem, n = 4, **** p < 0.0001, ** p < 0.01, one-way ANOVA with Dunnet’s multiple comparison test). **F**. Schematic representation of the genomic region corresponding to the Luc4 construct with pairs of CRISPR/Cas9 constructs designed to remove parts of the region. **G**. Ethidium bromide gel showing PCR products of the genomic region corresponding to the Luc4 construct. Shorter PCR products result from the removal of targeted genomic regions. **H**. SEMA7A mRNA expression in HUVEC infected with lentiviral constructs encoding Cas9 and gRNAs targeting genomic regions downstream of the SEMA7A gene and exposed to shear stress (18 dyn/cm^2^ for 6 h, data is shown as fold of control, mean ± sem, n = 3, * p < 0.05, repeated measures one-way ANOVA with Dunnet’s multiple comparison test). **I, J**. KLF2 (**I**) and SEMA7A (**J**) mRNA expression in HUVEC transfected with KLF2 or control siRNA (Neg-si) and exposed to shear stress (18 dyn/cm^2^ for 6 h, data are presented as fold of control, mean ± sem, n = 3, * p < 0.05, one-way ANOVA with Tukey’s multiple comparison test).

To test the function of KLF2 in regulating SEMA7A, we analyzed SEMA7A expression after KLF2 knockdown. We achieved a more than 50% knockdown of flow mediated KLF2 induction after siRNA transfection (Fig. 3I). This knockdown led to a strong decrease of SEMA7A expression in shear stress exposed ECs (Fig. 3J). Together, these results suggest that KLF2 mediates the observed shear stress regulation of SEMA7A expression.

Most reports focus on the positive effects enhancers have on gene expression. However, it is equally important that genes are being turned off. To validate a flow-responsive lost enhancer, we examined the known shear stress responsive gene CXCR4, which was strongly down-regulated upon exposure to shear stress (Fig. 1B, Suppl. Fig. 2B), which we confirmed by qPCR (Suppl. Fig. 5H). This gene was associated with a lost H3K27ac peak located 125 kb upstream of its TSS that overlapped with common ATAC peaks (Suppl. Fig. 5G). To confirm that this enhancer regulates CXCR4 expression, we used CRISPR/Cas9 to remove parts of the enhancer (Suppl. Fig. 5I, J). Removal of parts of the enhancer with CRISPR pairs 4+6 and 6+7 resulted in reduction of CXCR4 mRNA expression (Suppl. Fig. 5K), mimicking the shear stress response. We therefore identified and validated a shear stress responsive lost enhancer that regulates CXCR4 expression.

### Identification of flow-induced transcriptional program in zebrafish embryos

To study whether the transcriptional program we identified in adult human ECs is conserved during vascular development in other species, we compared gene expression in ECs in zebrafish embryos without blood flow with ECs from embryos experiencing normal blood flow. For this, we blocked blood flow in 48 hpf *Tg(kdrl:H2B-EGFP)^mu122^* embryos for 4 h and sorted ECs (Fig. 4A). RNA-Seq analysis revealed 1067 differentially expressed genes in ECs experiencing flow block compared to control zebrafish embryos. 584 genes were up- and 483 genes were downregulated (Fig. 4B, Suppl. Fig. 6A, B, Suppl. Data 1). We validated our RNA-Seq data by qPCR for select most up- and downregulated DE genes (Suppl. Fig. 6C), which showed a strong correlation between the two methods. GSEA pathway analysis revealed MAPK and Notch signaling as some of the most enriched pathways after flow block (Suppl Fig. 6D). Importantly, the orthologues of the genes that were among the most DE genes in the shear stress exposed HUVEC dataset were also regulated by blood flow in zebrafish embryos. Hence, *cxcr4a, kita, irs2a* and *irs2b* were upregulated, while *klf2a* and *sema7a* were downregulated by flow block (Fig. 4B). Expression changes were opposite to those we observed in HUVEC exposed to shear stress (Fig. 1B). Comparing the datasets with such opposite regulation in zebrafish and HUVEC revealed an 8 % overlap between the genes upregulated in zebrafish ECs after flow block and downregulated in HUVEC exposed to shear stress (Suppl. Fig. 6E). Conversely, there was a 6.2 % overlap of the genes downregulated in zebrafish ECs and upregulated in HUVEC (Suppl. Fig. 6E). This is a substantial overlap considering that only distinct EC populations in zebrafish embryos might show a flow response, which in addition might only occur at select developmental time points. Also, the experimental approach to interrogate for shear stress responses within ECs varied in these two systems (flow increase in the case of HUVEC, while we blocked blood flow in zebrafish embryos). Furthermore, the overlap was higher (13.7 %) when we took into consideration all DE genes (Fig. 4C, Suppl. Fig. 6E). This highlights the possibility that some genes might show differential flow responses, depending on the timing and the magnitude of shear stress. For example, ELMSAN1 is upregulated in HUVEC after 6 h of shear stress exposure and *elmsan1b* is also upregulated in zebrafish ECs after stopping blood flow. However, shorter shear stress exposure (30 min) resulted in downregulation of ELMSAN1 in HUVEC (Suppl. Fig. 1B) in line with our observations in zebrafish. DNA acetylation/de-acetylation will also require different sets of enzymes (histone acetylases and de-acetylases, respectively), which might not be available to the same extent in all cell populations.

**Figure 4.**
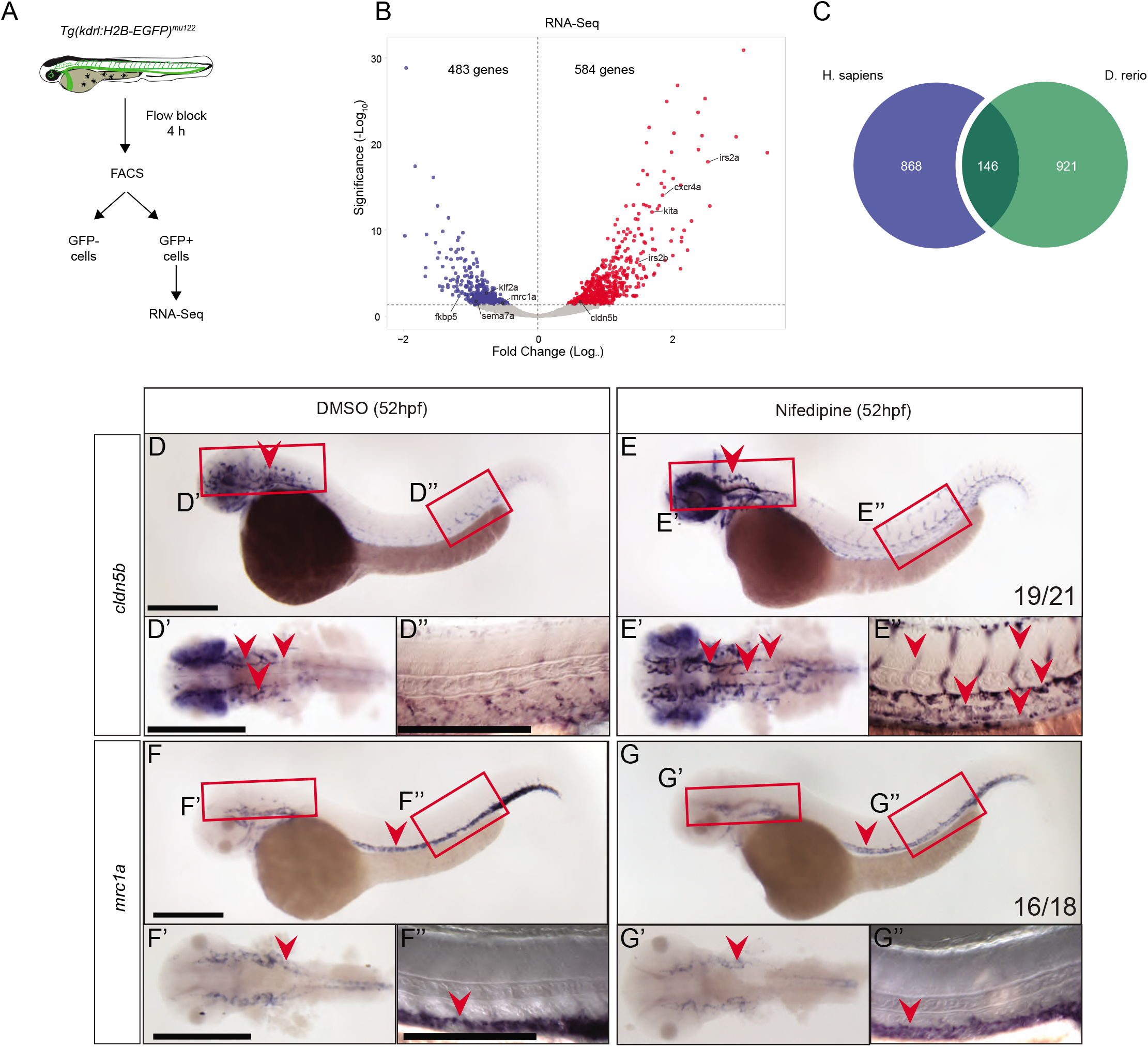
Blocking blood flow in zebrafish reveals genes regulated by shear stress in developing embryos. **A.** Schematic representation of EC FACS sorting from *Tg(kdrl:H2B-EGFP)^mu122^* embryos treated with nifedipine for 4 h at 48 hpf to induce flow block. Sorted GFP+ cells were used for RNA-Seq. As control, embryos were treated with a corresponding amount of DMSO. **B**. Volcano plot showing gene expression changes in ECs after flow block in zebrafish embryos. Differentially expressed (DE) genes (FDR < 0.05) are labelled in red (upregulated) and blue (downregulated after flow block). **C**. Overlap of flow-regulated genes in human and zebrafish datasets. All DE genes were used for this analysis. **D, E**. Whole mount in situ hybridization for *cldn5b* in 52 hpf embryos after 4 h of treatment with DMSO (**D**) or nifedipine (**E**). Regions in the head and the trunk highlighted with red boxes in **D** and **E** are magnified in **D’, E’** and **D”, E”**, respectively. Vascular expression is highlighted with red arrowheads. **F, G**. Whole mount in situ hybridization for *mrc1a* after flow block from 48 to 52 hpf. Regions in the head and the trunk highlighted with red boxes in **F** and **G** are magnified in **F’, G’** and **F”, G”**, respectively. Vascular expression is highlighted with red arrowheads. Numbers represent the number of embryos with the depicted expression pattern out of the total number of embryos analyzed. Scale bar is 300 μm.

We further validated vascular expression and flow responsiveness of *cldn5b* (Fig. 4D, E) and *mrc1a* (Fig. 4F, G) by whole mount in situ hybridization in zebrafish embryos. *cldn5b* was expressed in brain vasculature of 52 hpf embryos (Fig. 4D, arrowhead), while expression was weak in trunk vasculature (Fig. 4D, enlarged in D’ and D”). Flow block increased the expression of *cldn5b* in brain and trunk vasculature (Fig. 4E, arrowheads). This suggests that blood flow inhibits *cldn5b* expression, an observation that is in line with increased *cldn5b* expression after flow block in our RNA-Seq dataset (Fig. 4B). In contrast, expression of *mrc1a* in both brain and trunk vessels was reduced after flow block (Fig. 4F, G, enlarged in F’, F” and G’, G”). This correlated with *mrc1a* down-regulation by flow block in RNA-Seq (Fig. 4B) and reveals blood flow as a positive regulator of *mrc1a* expression. We furthermore confirmed upregulation of *adma* and *lyve1a* (Suppl. Fig. 7) and downregulation of *gpr182* and *id1* (Suppl. Fig. 8) upon flow block. These genes were not regulated in HUVEC but were differentially expressed on RNA-Seq of zebrafish ECs after flow block. In addition, we validated flow responsiveness of *sox18* and *klf2a* (Suppl. Fig. 9), which were regulated in zebrafish embryos and in HUVEC. In line with RNA-Seq results, flow block induced *sox18* expression in zebrafish blood vessels (Suppl. Fig. 9B, C). Interestingly, *sox18* was expressed in the dorsal aorta and posterior cardinal vein in 24 hpf embryos, but its expression was reduced in 36 and 48 hpf embryos, which experience increased blood flow (Suppl. Fig. Fig. 9A). Conversely, the expression of *klf2a* was higher in intersegmental blood vessels of 48 hpf embryos, compared to an earlier time point (36 hpf), which could be attributed to higher blood flow (Suppl. Fig. 9D). Blocking blood flow reduced intersegmental blood vessel expression of *klf2a* in accordance with our RNA-Seq results (Suppl. Fig. 9E, F).

## Discussion

It is widely appreciated that changes in hemodynamics have profound effects on EC biology. However, the underlying gene regulatory programs mediating these effects are poorly understood. In this report, we characterized the laminar shear stress response of HUVEC on the transcriptional and chromatin accessibility level. In addition, we provide evidence that reciprocal changes in gene expression occur in ECs of zebrafish embryos subjected to flow-block. Our analysis shows correlation between gene expression patterns and identified enhancers, with upregulated genes more commonly clustering with gained peaks, while downregulated genes associate with lost peaks. Thus, comparing ChIP-Seq and ATAC-Seq data from HUVEC revealed the presence of distinct sets of gene regulatory elements. We find a considerable overlap between gained H3K27ac ChIP-Seq and gained ATAC-Seq peaks. However, there is also an overlap between gained H3K27ac ChIP-Seq peaks and common ATAC-Seq peaks. This might suggest the existence of poised enhancers, which are positioned in regions of open chromatin, but have not obtained H3K27ac marks^38, 65^.

We find that shear stress upregulates the expression of the transcription factors KLF2 and KLF4, which previous studies identified as master regulators of the endothelial shear stress response^5, 66^. We find KLF2 and KLF4 upregulation already after 30 minutes of shear stress exposure, identifying these genes being among the earliest transcriptional responses to shear stress^56^. Loss of KLF2 has been reported to reduce vascular smooth muscle cell and pericyte coverage^67, 68^, lead to high cardiac output failure^69^ and cardiac valve defects in zebrafish^70^ and mouse^71^. High *klf2a* expression in developing veins in zebrafish is needed to restrict smooth muscle cells to the dorsal aorta^72^. Further studies showed that *klf2a* controls VEGF signaling during aortic arch development^73^. In addition to these requirements of KLF2 during embryonic development, KLF2 also regulates vascular tone^74^, represses inflammatory responses^75^ and induces endothelial quiescence and atheroprotection^76^. Studies analyzing the human disease condition cerebral cavernous malformations (CCM) also showed that increases in KLF2 mRNA levels can contribute to disease progression^77^, illustrating the need of a tight control of KLF2 expression. Given the multitude of processes KLF2 functions in, additional work is needed to separate primary from secondary effects within the vasculature. Our work provides an inroad to answering these questions through the identification of KLF binding sites in regions of the chromatin that show marks of active enhancers upon flow exposure. Functionally interrogating these regions will help in delineating direct KLF2 target genes in addition to other transcription factors which might bind to the novel motifs we identified.

We initiated these studies by analyzing putative enhancer elements driving the expression of SEMA7A. We chose SEMA7A due to its regulation by shear stress and reported role in blood vessel homeostasis^78^. Our bioinformatic analysis showed the presence of putative enhancer elements about 50 KB downstream of the SEMA7A TSS. We confirmed the flow response of this enhancer element using luciferase reporter constructs and through deleting it from the HUVEC genome. Therefore, our approach can lead to the identification of shear stress responsive enhancer elements. However, about 80% of the enhancers we tested did not change the expression of luciferase reporter constructs. This is within the predicted range^62^. Several reasons can account for the lack of reporter activity. Enhancers often function together with the promoters of the genes they regulate^64, 79^. Thus, using a gene’s endogenous promoter instead of a generic minimal promoter might increase the number of enhancers functioning in reporter assays. The genomic context of a given enhancer might also influence its activity^80^. In addition, a growing body of evidence suggests that genes are being regulated through the combinatorial action of several “weak” enhancers^63, 81^. In the future, interrogating these interactions within our dataset will likely contribute significant insights into shear stress mediated gene regulation. It will also be of importance to understand the effects different flow regimes exert on endothelial cells. We previously characterized enhancer elements and transcription factors in ECs exposed to oscillatory shear stress^33^, while He et al. generated a similar dataset for ECs exposed to pulsatile shear stress^34^. Furthermore, Andueza et al. generated datasets for disturbed flow^32^. Comparing these data to our data investigating laminar shear stress responses will help in elucidating how ECs might adapt to changes in flow during embryonic development or disease settings.

Our data significantly expand previous studies that focused more on the activating effects of shear stress on gene expression and the transcription factors mediating these effects^32^. In this study, we also analyzed lost enhancers upon flow exposure, which might be equally important for regulating vascular morphogenesis. For instance, during angiogenesis in zebrafish, expression of *flt4* and *cxcr4a* needs to be downregulated in order to prevent ectopic blood vessel sprouting^82–85^, while prolonged ETV2 expression during embryogenesis can lead to vascular defects^86^. Our data now provide a resource to investigate the influence of lost enhancers on endothelial gene expression. An unexpected finding was the enrichment of ETV2/ETS binding sites in this enhancer subset. During embryogenesis ETV2 is required for EC differentiation, but is strongly downregulated at later stages of blood vessel development^87–91^. In undifferentiated progenitor cells in culture, ETV2 expression is needed to commit these cells to the endothelial lineage^92^, while overexpression in HUVECs allows these cells to obtain a more embryonic EC state^93^. This overexpression leads to an upregulation of KLF2 and KLF4 expression with ETV2 being bound to KLF2 and KLF4 TSS. By contrast, continuous ETV2 expression in mouse embryos reduces KLF2 and KLF4 levels and leads to vascular anomalies^86^. Of note, analysis of several vein specific enhancers revealed the presence of multiple ETS binding sites. Reporter constructs using these enhancers displayed robust expression during early embryonic stages but were downregulated after E13 in mice and did not show expression in the mature microvasculature^94^. Together, these results suggest that suppression of ETV2 and ETV2 target gene expression might be required for proper shear stress responses and that expression of KLF2 and KLF4 might initially rely on ETV2, but that continuous expression requires ETV2 downregulation. Our results showing a loss of binding sites for ETS and FLI transcription factors in response to shear stress support this notion and will enable the discovery of further ETV2 target genes. How KLF and ETS TFs function in concert and the mechanisms controlling their regulatory interdependence will provide important future insights into shear stress mediated endothelial differentiation and vascular maturation.

Zebrafish have advanced as a widely used model system to study cardiovascular development and disease^95^. Our findings that many genes are similarly regulated by shear stress when compared to HUVEC underscore the conserved nature of shear stress responses. Importantly, zebrafish can be used to investigate tissue and temporal differences in these responses. Future studies will be needed to reveal how gene regulatory circuits change during blood vessel development and decipher the contribution shear stress has on these processes and thereby vascular morphogenesis.

## Author contributions

R.T. and A.F.S. conceived the experiments. A.F.S., H.S., J.M.V. and Z.A. supervised the work. R.T. performed cell culture experiments and performed data analysis. E.V.L. performed zebrafish experiments. N.D. and N.Y. performed data analysis. M.O.-S., O.B. and R.T. performed shear stress experiments. R.T. and A.F.S. wrote the manuscript. All authors provided manuscript input and discussed and commented on the work.

## Acknowledgements

We would like to thank Gerd Blobel for critical reading of the manuscript. This work was supported from funds provided by the Max-Planck society (A.F.S.), Deutsche Forschungsgemeinschaft (DFG) grant SI 1374/4-1, SI 1374/5-1, SI 1374/ 3-2 (A.F.S.) and the Perelman School of Medicine and the Cardiovascular Research Institute of the University of Pennsylvania (A.F.S.).

---

**Supplementary Figure 1. Exposing HUVEC to shear stress for 30 minutes regulates the expression of a distinct set of genes. A**. Volcano plot representing differentially expressed (DE) genes (FDR < 0.05) upon HUVEC exposure to shear stress of 18 dyn/cm^2^ for 30 min. Upregulated genes are labelled in red and downregulated genes in blue. **B, C**. Top 20 upregulated genes (**B**) and 17 genes downregulated in HUVEC exposed to shear stress for 30 minutes (**C**).

**Supplementary Figure 2. Overlap of shear stress regulated genes between different studies. A, B**. 20 genes most up- (**A**) or downregulated (**B**) upon 6 h exposure to 18 dyn/cm^2^ shear stress (RNA-Seq results). **C**. Correlation of mRNA expression measured by RNA-Seq and qPCR in shear stress exposed HUVEC (n =2). **D**. Over-representation (ORA) analysis of KEGG pathways regulated in the RNA-Seq dataset. **E**. Comparison of DE genes regulated by shear stress in this study with published datasets. RNA-Seq data of Maleszewska et al. (HUVEC exposed to 20 dyn/cm^2^ for 72 h), microarray data of Dekker et al. (HUVEC with adenoviral overexpression of KLF2 for 7 days) and metaanalysis of multiple microarray studies (Maimari et al.) are listed. The amount of up- and down-regulated genes overlapping with our data is presented. The total amount of overlapping genes (irrespective of up- or down-regulation) is additionally shown as percentage compared to the total number of DE genes in our RNA-Seq.

**Supplementary Figure 3. Verification of H3K27ac ChIP-Seq data. A.** Pairwise correlation analysis of ChIP-Seq HUVEC samples (cultured under static, “Control”, or shear stress conditions, “Flow”) with HUVEC H3K27ac ChIP-Seq from ENCODE. Pearson coefficient values for each comparison are represented. **B**. Verification of selected gained, lost and common H3K27ac ChIP-Seq peaks with ChIP-qPCR. Additionally, a region with no H3K27ac ChIP-Seq enrichment was tested as negative control. The data are presented as % of input (mean + sem, n = 3, * p < 0.05 compared to corresponding control, two-tailed paired t-test).

**Supplementary Figure 4. Comparison of ATAC-Seq and H3K27ac ChIP-Seq data sets and DNA motif analysis. A.** Annotation of all gained, lost and common ATAC-Seq peaks to the defined regions of the genome. **B**. Overlap of gained and lost ATAC-Seq peaks with gained, lost and common H3K27ac ChIP-Seq peaks. Numbers represent percentage of overlap. **C.** Overlap of gained and lost H3K27ac ChIP-Seq peaks with gained, lost and common ATAC-Seq peaks. Numbers represent percentage of overlap. **D, E**. de novo motif analysis of DNA motifs enriched in gained (**D**) or lost (**E**) ATAC-Seq peaks that overlap with gained and lost H3K27ac ChIP-Seq peaks, respectively. Most enriched motifs (top 10 for gained and all 7 for lost) and corresponding best match are shown. **F, G**. Known motifs enriched in common ATAC-Seq peaks that overlap with gained (**F**) or lost (**G**) H3K27ac-ChIP-Seq peaks. The 10 most enriched motifs are shown.

**Supplementary Figure 5. Functional analysis of flow responsive enhancers. A.** Schematic representation of luciferase constructs with a putative enhancer (E) upstream of a minimal promoter (minP) and firefly luciferase gene (Luc). **B**. Luciferase assays with constructs carrying putative enhancers associated with DE genes in HUVEC exposed to 18 dyn/cm^2^ for 24 h (data is presented as arbitrary units (a.u.) after normalization to the signal of renilla luciferase; mean ± sem, n = 3 - 5, * p < 0.05, **** p < 0.001, compared to a corresponding static control, two-tailed paired t-test). **C**. Schematic representation of dCas9-p300 construct designed to target putative enhancers to affect gene expression. **D**. Schematic representation of the removal of a putative enhancer (E) with Cas9 and two specific gRNAs (P, promoter; ORF, open reading frame). **E**. Luciferase assay showing luciferase activity of Luc2 and Luc4 constructs, as well as a construct containing both regions (Luc2+4) in HUVEC exposed to 18 dyn/cm^2^ for 24 h (the data is presented as arbitrary units (a.u.) after normalization to the signal of renilla luciferase; mean ± sem, n = 3, * p < 0.05, two-tailed paired t-test). **F.** Luciferase activity of Luc4 construct targeted by dCas9-p300 and specific gRNAs in HEK293T17 cells (the data are presented as arbitrary units (a.u.) after normalization to the signal of a renilla luciferase construct; mean ± sem, n = 3, * p < 0.05, ** p < 0.01, one-way ANOVA with Dunnet’s multiple comparison test). **G**. Normalized ATAC-Seq and H3K27ac ChIP-Seq tracks around the CXCR4 gene. The bars below ATAC-Seq and H3K27ac ChIP-Seq tracks represent corresponding common (grey) or lost (blue) peaks. **H**. CXCR4 mRNA expression in HUVEC exposed to 18 dyn/cm^2^ for 6 h (Log2 fold of control, mean ± sem, n = 3, * p < 0.05, two-tailed paired t-test). **I**. Schematic representation of the genomic region upstream of the CXCR4 gene and the pairs of CRISPR/Cas9 constructs designed to remove parts of the region. **J**. Ethidium bromide gel confirming removal of targeted genomic regions. PCR products indicated with an asterisk are shorter than in corresponding controls (Co) and result from successful CRISPR targeting (Cr). **K**. CXCR4 mRNA expression in HUVEC infected with lentiviral constructs encoding Cas9 and gRNAs targeting genomic regions upstream of the CXCR4 gene (data are shown as fold of control, mean ± sem, n = 4, ** p < 0.01, repeated measures one-way ANOVA with Dunnet’s multiple comparison test).

**Supplementary Figure 6. Analysis of flow regulated genes in zebrafish. A.** 20 genes most up- (**A**) or downregulated (**B**) in zebrafish ECs 4 h after blocking blood flow (48 - 52 hpf, RNA-Seq results). **C**. Correlation of mRNA expression measured by RNA-Seq and qPCR in zebrafish endothelial cells after blocking blood flow (n =2). **D**. Gene set enrichment analysis (GSEA) of KEGG pathways regulated in the zebrafish RNA-Seq dataset. **E**. Overlap of up- and downregulated, as well as all DE genes in zebrafish with human orthologues regulated by shear stress.

**Supplementary Figure 7. Analysis of genes downregulated by flow. A.** Whole mount in situ hybridization for *adma* shows no detectable vascular expression in 24, 36 and 48 hpf zebrafish embryos. **B, C**. Nifedipine treatment from 48 to 52 hpf induces *adma* expression in brain blood vessels (black arrowheads) **C**, enlarged in **C’**), while expression is undetectable in DMSO controls (**B**, enlarged in **B’**). **D**. Whole mount hybridization for *lyve1a* showing no vascular expression in 24, 36 and 48 hpf embryos. **E, F**. *lyve1a* expression in 52 hpf embryos after 4 h of treatment with DMSO (**E**) or nifedipine (**F**). Enlarged images of the head (**E’** and **F’**) and the trunk (**E’**’ and **F”**) regions highlighted by red rectangles in **E** and **F**. Flow block induces an increased *lyve1a* expression in the common cardinal vein (white arrowheads in **E**’, **F** and **F’**) and in caudal plexus (blue arrowheads in **F** and **F”**). Numbers represent number of embryos with depicted expression pattern out of the total number of embryos tested. Scale bar is 300 μm.

**Supplementary Figure 8. Analysis of genes upregulated by flow. A.** Whole mount in situ hybridization for *grp182* in 24, 36 and 48 hpf zebrafish embryos. *grp182* is expressed in trunk axial vasculature (black box) and in the primordial midbrain channel (black arrowhead) at 24 hpf and in the dorsal aorta (red arrowheads) and posterior cardinal vein (blue arrowheads) at 36 and 48 hpf. **B, C**. *grp182* expression in 52 hpf embryos after 4 h treatment with DMSO (**B**) or nifedipine (**C**). Enlarged images of the head (**B’** and **C’**) and the trunk (**B’**’ and **C”**) regions highlighted by red rectangles in **B** and **C**. Flow block decreases *grp182* expression in lateral dorsal aorta (red arrowheads) and posterior cardinal vein (blue arrowheads). **D**. Whole mount in situ hybridization for *id1* in 24, 36 and 48 hpf zebrafish embryos. Expression of *id1* is detected in the neural tube (blue arrowheads), hypochord (red arrowheads), axial vasculature (black box) and intersegmental blood vessels (black arrowheads). **E, F**. *id1* expression in 52 hpf embryos after 4 h treatment with DMSO (**E**) or nifedipine (**F**). Enlarged images of the head (**E’** and **F’**) and the trunk (**E’**’ and **F”**) regions highlighted by red and black rectangles in **E** and **F**. Flow block reduces *id1* expression in intersegmental vessels (black arrowheads) and in axial vessels (white brackets), while expression in the neural tube (blue arrowheads) and hypochord (red arrowheads) remains unaffected. Numbers represent number of embryos with depicted expression pattern out of the total number of embryos tested. Scale bar is 300 μm.

**Supplementary Figure 9. Regulation of *sox18* and *klf2a* expression by flow in zebrafish embryos. A**. Whole mount in situ hybridization for *sox18* in 24, 36 and 48 hpf zebrafish embryos. The black box highlights vascular expression in the trunk at 24 hpf, which is reduced at later stages. **B, C**. *sox18* expression in 52 hpf embryos after 4 h treatment with DMSO (**B**) or nifedipine (**C**). Enlarged images of the head (**B’** and **C’**) and the trunk (**B’**’ and **C”**) regions highlighted by red rectangles in **B** and **C**. Black arrowheads indicate vascular expression, which is increased after inhibition of blood flow. **D**. Whole mount in situ hybridization for *klf2a* in 24, 36 and 48 hpf zebrafish embryos. Black arrowheads indicate expression in trunk axial vessels at 24 and 36 hpf and the black box highlights increased expression in axial vessels, as well as in intersegmental vessels at 48 hpf. **E, F**. *klf2a* expression in 52 hpf embryos after 4 h treatment with DMSO (**E**) or nifedipine (**F**). Enlarged images of the head (**E**’ and **F**’) and the trunk (**E”** and **F”**) regions highlighted by red boxes in **E** and **F**. Black arrowheads indicate vascular expression, which is decreased after flow block in the brain and in the trunk. Numbers represent number of embryos with depicted expression pattern out of the total number of embryos analyzed. Scale bar is 300 μm.

