## Supplementary Figures for "DNA motif analysis of shear stress responsive endothelial enhancers reveals differential association of KLF and ETV/ETS binding sites with gained and lost enhancers"

A

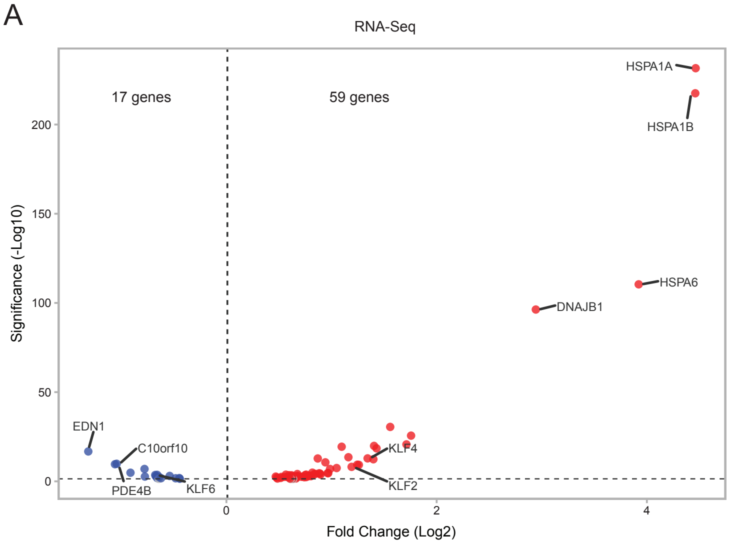


B

| ENSEMBL Gene ID | log2FoldChar | lfcSE | padj | Gene Name |
|---|---|---|---|---|
| ENSG00000204389 | 4.46393816 | 0.13601455 | 2.61E-232 | HSPA1A |
| ENSG00000204388 | 4.46031914 | 0.14027444 | 3.00E-218 | HSPA1B |
| ENSG00000173110 | 3.92204737 | 0.17237454 | 3.77E-111 | HSPA6 |
| ENSG00000132002 | 2.94345393 | 0.13841216 | 4.96E-97 | DNAJB1 |
| ENSG00000013441 | 1.75667656 | 0.1557838 | 2.42E-26 | CLK1 |
| ENSG00000117616 | 1.71267223 | 0.16759484 | 1.96E-21 | RSRP1 |
| ENSG00000120694 | 1.55819638 | 0.12728521 | 3.14E-31 | HSPH1 |
| ENSG00000124216 | 1.42812358 | 0.1475603 | 3.15E-19 | SNAI1 |
| ENSG00000151929 | 1.40452248 | 0.1402296 | 1.37E-20 | BAG3 |
| ENSG00000188483 | 1.39806471 | 0.17346433 | 4.31E-13 | IER5L |
| ENSG00000136826 | 1.34308432 | 0.1634457 | 1.35E-13 | KLF4 |
| ENSG00000160570 | 1.26020551 | 0.17783703 | 5.81E-10 | DEDD2 |
| ENSG00000170345 | 1.24456699 | 0.17488521 | 4.92E-10 | FOS |
| ENSG00000127528 | 1.19094644 | 0.17743925 | 7.73E-09 | KLF2 |
| ENSG00000004478 | 1.1605387 | 0.13809586 | 3.04E-14 | FKBP4 |
| ENSG00000080824 | 1.09653281 | 0.11079645 | 4.03E-20 | HSP90AA1 |
| ENSG00000204569 | 1.04857835 | 0.16180978 | 3.51E-08 | PPP1R10 |
| ENSG00000197019 | 0.98590745 | 0.15698868 | 1.19E-07 | SERTAD1 |
| ENSG00000140403 | 0.96888764 | 0.18088438 | 2.56E-05 | DNAJA4 |
| ENSG00000198576 | 0.96312538 | 0.1795659 | 2.55E-05 | ARC |

C

| ENSEMBL Gene ID | log2FoldChar | lfcSE | padj | Gene Name |
|---|---|---|---|---|
| ENSG00000078401 | -1.3139386 | 0.14209871 | 1.78E-17 | EDN1 |
| ENSG00000184588 | -1.0601802 | 0.14694164 | 2.53E-10 | PDE4B |
| ENSG00000165507 | -1.0438904 | 0.14302887 | 1.45E-10 | C10orf10 |
| ENSG00000137331 | -0.9125793 | 0.16691337 | 1.48E-05 | IER3 |
| ENSG00000139318 | -0.7791677 | 0.12388776 | 1.17E-07 | DUSP6 |
| ENSG00000116717 | -0.7754477 | 0.17784408 | 0.002150288 | GADD45A |
| ENSG00000067082 | -0.6769533 | 0.13959285 | 0.000274993 | KLF6 |
| ENSG00000156030 | -0.6722485 | 0.14918951 | 0.001220766 | ELMSAN1 |
| ENSG00000124575 | -0.6585195 | 0.1700084 | 0.014378774 | HIST1H1D |
| ENSG00000118515 | -0.6583668 | 0.13477376 | 0.000236013 | SGK1 |
| ENSG00000177352 | -0.6435607 | 0.18037964 | 0.041628795 | CCDC71 |
| ENSG00000113369 | -0.6403809 | 0.15122304 | 0.003450353 | ARRDC3 |
| ENSG00000143067 | -0.6215881 | 0.16299926 | 0.017798993 | ZNF697 |
| ENSG00000163788 | -0.5397281 | 0.11818853 | 0.000950745 | SNRK |
| ENSG00000185112 | -0.4830494 | 0.12738184 | 0.019104008 | FAM43A |
| ENSG00000143878 | -0.448177 | 0.11611076 | 0.014962019 | RHOB |
| ENSG00000124766 | -0.4432221 | 0.12282025 | 0.036081265 | SOX4 |

Tsaryk et al., Supplementary Fig. 1

A

| ENSEMBL Gene ID | log2FoldChar | lfcSE | padj | Gene Name |
|---|---|---|---|---|
| ENSG00000164530 | 7.23157246 | 0.5280092 | 1.17E-39 | PI16 |
| ENSG00000104783 | 6.32277206 | 0.57912565 | 5.64E-25 | KCNN4 |
| ENSG00000136999 | 5.85304792 | 0.41122402 | 9.33E-43 | NOV |
| ENSG00000154734 | 5.75946696 | 0.36967531 | 4.36E-51 | ADAMTS1 |
| ENSG00000159871 | 5.70060654 | 0.52796033 | 1.93E-24 | LYPD5 |
| ENSG00000174600 | 5.59695605 | 0.3902341 | 2.59E-43 | CMKLR1 |
| ENSG00000165272 | 5.55498339 | 0.4965064 | 3.05E-26 | AQP3 |
| ENSG00000138061 | 5.35585517 | 0.37750583 | 1.59E-42 | CYP1B1 |
| ENSG00000095752 | 5.20564682 | 0.55119064 | 1.27E-18 | IL11 |
| ENSG00000127528 | 5.18998818 | 0.32922041 | 3.58E-52 | KLF2 |
| ENSG00000136826 | 5.18397413 | 0.31076796 | 2.35E-58 | KLF4 |
| ENSG00000138623 | 5.10866282 | 0.36068163 | 2.01E-42 | SEMA7A |
| ENSG00000115461 | 4.81144856 | 0.57753478 | 1.69E-14 | IGFBP5 |
| ENSG00000138670 | 4.66333256 | 0.45288322 | 3.40E-22 | RASGEF1B |
| ENSG00000156427 | 4.59630516 | 0.61176361 | 9.00E-12 | FGF18 |
| ENSG00000163273 | 4.54553639 | 0.6422798 | 1.97E-10 | NPPC |
| ENSG00000159167 | 4.51412326 | 0.64154326 | 2.56E-10 | STC1 |
| ENSG00000103196 | 4.44455529 | 0.50811566 | 5.73E-16 | CRISPLD2 |
| ENSG00000158955 | 4.35011621 | 0.57866494 | 8.81E-12 | WNT9B |
| ENSG00000136153 | 4.20294978 | 0.30668654 | 1.14E-39 | LMO7 |

B

| ENSEMBL Gene ID | log2FoldChar | lfcSE | padj | Gene Name |
|---|---|---|---|---|
| ENSG00000120738 | -4.4450289 | 0.50319511 | 2.71E-16 | EGR1 |
| ENSG00000131386 | -4.295856 | 0.44044927 | 6.88E-20 | GALNT15 |
| ENSG00000157404 | -4.1179487 | 0.36338554 | 6.98E-27 | KIT |
| ENSG00000165507 | -4.0506711 | 0.34476833 | 6.69E-29 | C10orf10 |
| ENSG00000091879 | -3.8945261 | 0.33828627 | 9.34E-28 | ANGPT2 |
| ENSG00000185950 | -3.5480676 | 0.58826512 | 1.22E-07 | IRS2 |
| ENSG00000121966 | -3.4732088 | 0.64951835 | 4.69E-06 | CXCR4 |
| ENSG00000088756 | -3.4616065 | 0.32997433 | 4.63E-23 | ARHGAP28 |
| ENSG00000185477 | -3.3086477 | 0.63290089 | 8.26E-06 | GPRIN3 |
| ENSG00000196421 | -3.1331681 | 0.52521289 | 1.72E-07 | LINC00176 |
| ENSG00000175746 | -3.0777502 | 0.39842478 | 1.91E-12 | C15orf54 |
| ENSG00000118407 | -3.0379694 | 0.64954491 | 0.000106491 | FILIP1 |
| ENSG00000171408 | -3.0088392 | 0.36997637 | 8.22E-14 | PDE7B |
| ENSG00000198355 | -2.9427072 | 0.31875833 | 7.75E-18 | PIM3 |
| ENSG00000005187 | -2.9318501 | 0.36192126 | 1.05E-13 | ACSM3 |
| ENSG00000128594 | -2.8804755 | 0.55544442 | 1.02E-05 | LRRC4 |
| ENSG00000137266 | -2.8721365 | 0.38840704 | 2.11E-11 | SLC22A23 |
| ENSG00000151967 | -2.8712331 | 0.44830357 | 1.38E-08 | SCHIP1 |
| ENSG00000004799 | -2.8600017 | 0.45302171 | 2.42E-08 | PDK4 |
| ENSG00000152207 | -2.8474613 | 0.70002554 | 0.001222303 | CYSLTR2 |

C

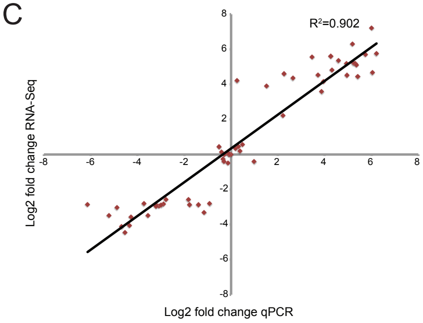


E

| Genes | Our data | Maleszewska et al | | Maimari et al | | Dekker et al | |
|---|---|---|---|---|---|---|---|
| | DE genes | DE genes | Overlap | DE genes | Overlap | DE genes | Overlap |
| Upregulated | 647 | 615 | 233 | 1003 | 249 | 600 | 111 |
| Downregulated | 367 | 835 | 104 | 635 | 71 | 439 | 44 |
| Total | 1014 | 1450 | 350 | 1638 | 330 | 1039 | 163 |
| % | | | 34.5% | | 32.5% | | 15.7% |

D

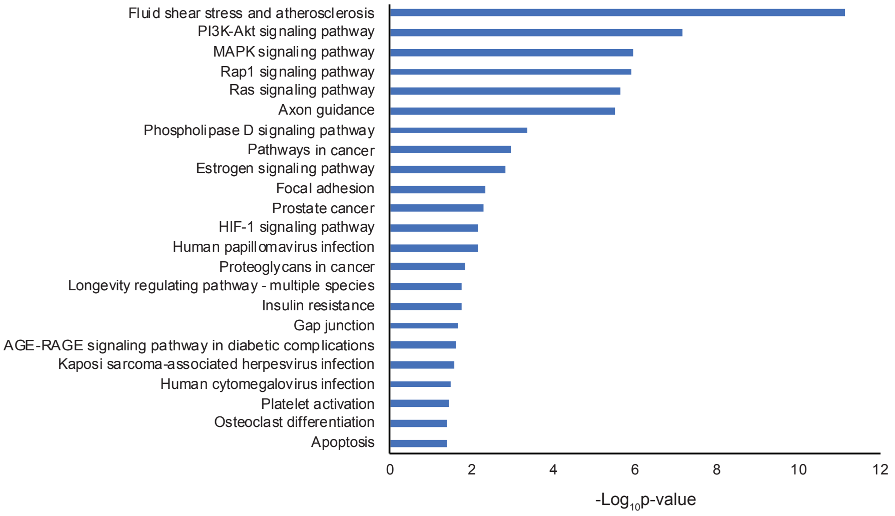


Tsaryk et al., Supplementary Fig. 2

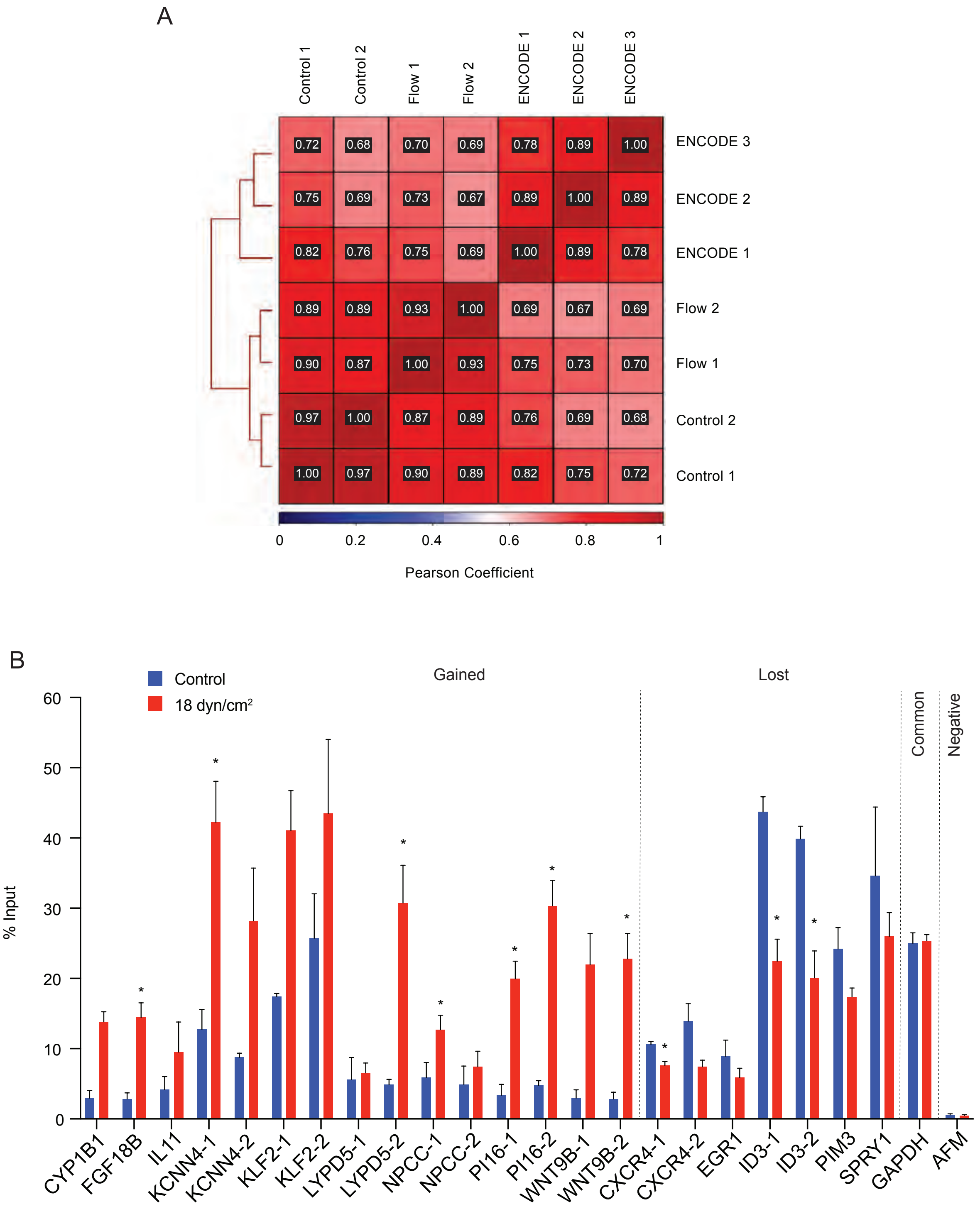


Tsaryk et al., Supplementary Fig. 3

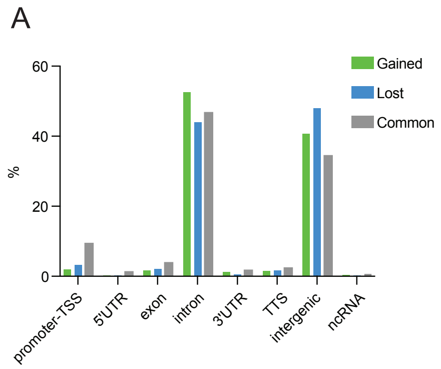


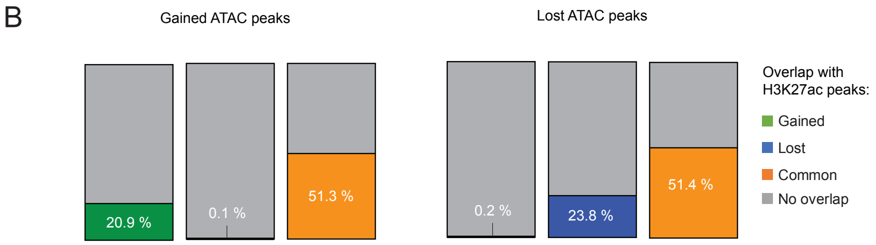


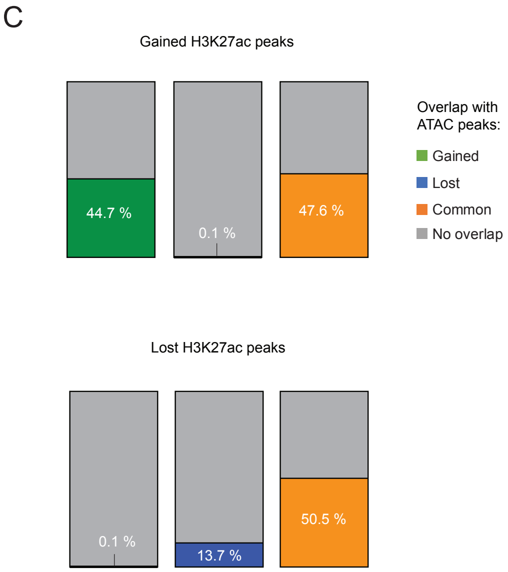


## D

Novel motifs in Gained ATAC/Gained H3K27ac peaks

| Motif | Best Match | Log P-value |
|---|---|---|
| TGGGTGTGGC | Klf1/MA0493.1/Jaspar(0.933) | -2.83E+02 |
| CCCTTCCTGC | ETS1(ETS)/Jurkat-ETS1-ChIP-Seq(GSE17954)/Homer(0.815) | -5.56E+01 |
| GATTTGACACAC | BATF::JUN/MA0462.1/Jaspar(0.606) | -4.63E+01 |
| GAAGTCTGGATG | PB0060.1_Smad3_1/Jaspar(0.625) | -4.37E+01 |
| TGACTCAC | Fra1(bZIP)/BT549-Fra1-ChIP-Seq(GSE46166)/Homer(0.860) | -4.26E+01 |
| CAGTATCAATCA | PB0144.1_Lef1_2/Jaspar(0.798) | -4.24E+01 |
| TCATGACAGA | MAFG::NFE2L1/MA0089.1/Jaspar(0.723) | -3.73E+01 |
| GGAAGGGGAGGA | SP2/MA0516.1/Jaspar(0.679) | -3.62E+01 |
| AGACATCAGC | c-Jun-CRE(bZIP)/K562-cJun-ChIP-Seq(GSE31477)/Homer(0.699) | -3.52E+01 |
| AGAGGTTATT | COUP-TFII(NR)/Artia-Nr2f2-ChIP-Seq(GSE46497)/Homer(0.692) | -3.48E+01 |

## E

Novel motifs in Lost ATAC/Lost H3K27ac peaks

| Motif | Best Match | Log P-value |
|---|---|---|
| TTCCTGTTTA | Ets1-distal(ETS)/CD4+-PolII-ChIP-Seq(Barski_et_al.)/Homer(0.852) | -7.33E+01 |
| AAAACAAT | Sox5/MA0087.1/Jaspar(0.887) | -3.81E+01 |
| ATAAACAGCCTT | Foxf1(Forkhead)/Lung-Foxf1-ChIP-Seq(GSE77951)/Homer(0.637) | -3.72E+01 |
| ACTTTTTCTTTG | PB0093.1_Zfp105_1/Jaspar(0.744) | -3.67E+01 |
| AAACATCAGG | ZBTB18/MA0698.1/Jaspar(0.758) | -3.51E+01 |
| TGATCCCCAAGA | MZF1/MA0056.1/Jaspar(0.671) | -2.88E+01 |
| AAGATTATGTTT | PB0016.1_Foxj1_1/Jaspar(0.671) | -2.81E+01 |

## F

Known motifs in Common ATAC/Gained H3K27ac peaks

| Motif | Motif Name | Log P-value |
|---|---|---|
| CTGGGTGTGGCC | EKLF(Zf)/Erythrocyte-Klf1-ChIP-Seq(GSE20478) | -1.24E+01 |
| TTGACCTTTG | RARa(NR)/K562-RARa-ChIP-Seq(Encode) | -1.22E+01 |
| GATGAGTCATCC | Fosl2(bZIP)/3T3L1-Fosl2-ChIP-Seq(GSE56872) | -1.14E+01 |
| AAGGTCACCGTGACC | ERE(NR),IR3/MCF7-ERa-ChIP-Seq | -1.14E+01 |
| GATGACTCAT | JunB(bZIP)/DendriticCells-Junb-ChIP-Seq(GSE36099) | -1.08E+01 |
| GCCACACCCA | Klf4(Zf)/mES-Klf4-ChIP-Seq(GSE11431) | -1.05E+01 |
| CCACTTCCTGTT | Etv2(ETS)/ES-ER71-ChIP-Seq(GSE59402) | -1.01E+01 |
| GGTCAAAGGTCA | COUP-TFII(NR)/K562-NR2F1-ChIP-Seq(Encode) | -1.01E+01 |
| AGGGTGTGGC | KLF5(Zf)/LoVo-KLF5-ChIP-Seq(GSE49402) | -1.00E+01 |
| GGGGGGGG | Maz(Zf)/HepG2-Maz-ChIP-Seq(GSE31477) | -0.98E+00 |

## G

Known motifs in Common ATAC/Lost H3K27ac peaks

| Motif | Motif Name | Log P-value |
|---|---|---|
| ATTTCCTGTTTT | ETV1(ETS)/GIST48-ETV1-ChIP-Seq(GSE22441) | -4.96E+01 |
| AAACAACG | Ets1-distal(ETS)/CD4+-PolII-ChIP-Seq(Barski_et_al.) | -4.50E+01 |
| CCCTAACTTACT | ERG(ETS)/VCaP-ERG-ChIP-Seq(GSE14097) | -4.30E+01 |
| CAGTGGAAAT | GABPA(ETS)/Jurkat-GABPa-ChIP-Seq(GSE17954) | -3.91E+01 |
| CAAGGGCACT | SPDEF(ETS)/VCaP-SPDEF-ChIP-Seq(SRA014231) | -3.73E+01 |
| CTATAAACTCTG | ETS1(ETS)/Jurkat-ETS1-ChIP-Seq(GSE17954) | -3.54E+01 |
| TGGCATGTTGCA | Etv2(ETS)/ES-ER71-ChIP-Seq(GSE59402) | -3.36E+01 |
| TTTCTCCACGCA | ETV4(ETS)/HepG2-ETV4-ChIP-Seq(ENCODE) | -2.65E+01 |
| CCTGAAATTGTT | EWS:ERG-fusion(ETS)/CADO_ES1-EWS:ERG-ChIP-Seq(SRA014231)/ | -2.61E+01 |
| AGTCTGGGATTC | Fli1(ETS)/CD8-FLI-ChIP-Seq(GSE20898) | -2.43E+01 |

Tsaryk et al., Supplementary Fig. 4

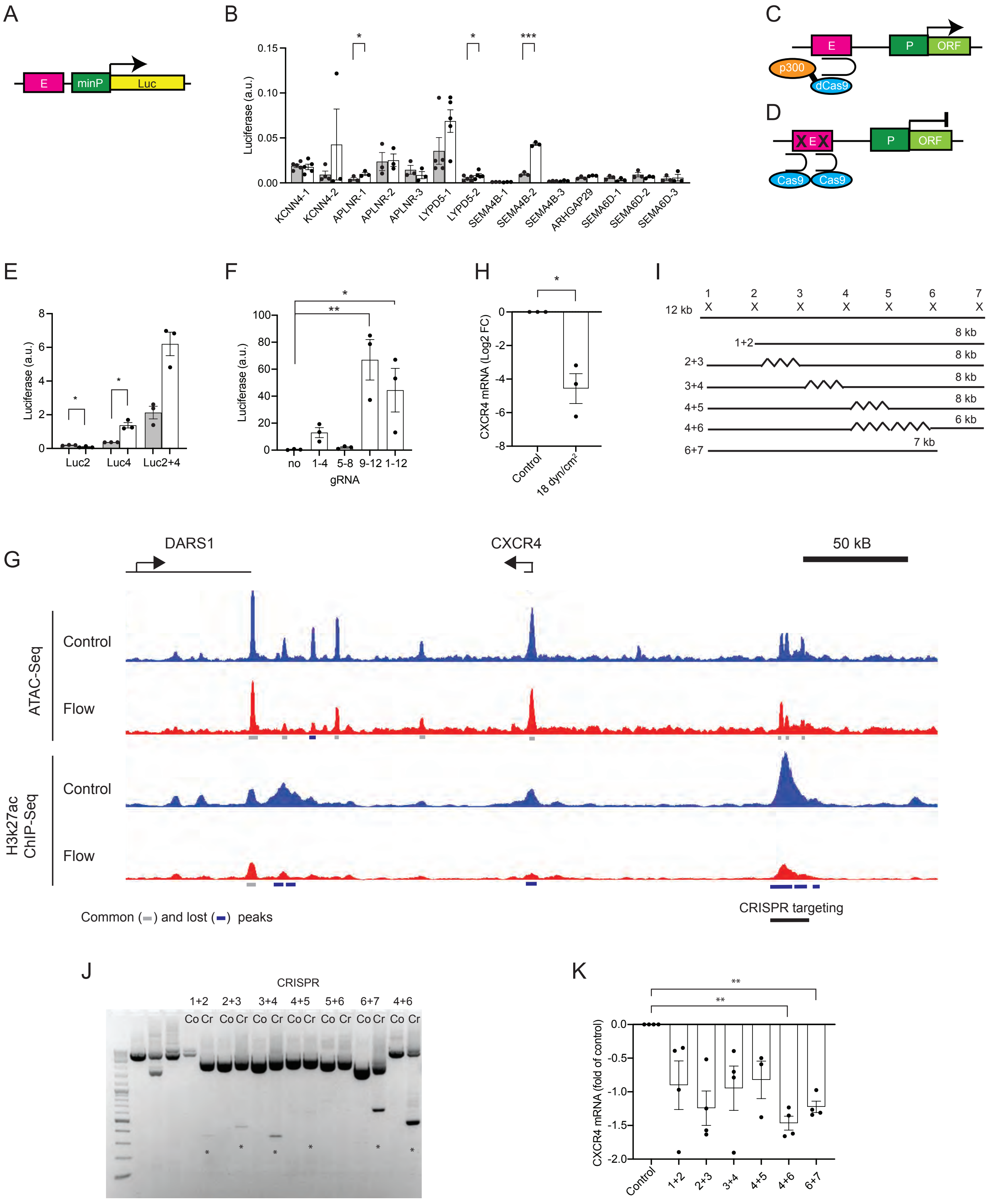


Tsaryk et al., Supplementary Fig. 5

A

| ENSEMBL Gene ID | log2FoldChar | lfcSE | padj | gene_name |
|---|---|---|---|---|
| ENSDARG00000052646 | 3.42153839 | 0.35031736 | 1.03E-19 | ftr69 |
| ENSDARG00000039943 | 3.06727405 | 0.24633644 | 1.27E-31 | fam46ba |
| ENSDARG00000098155 | 2.95610381 | 0.28914694 | 1.44E-21 | si:dkeyp-3b12.11 |
| ENSDARG00000019874 | 2.56354094 | 0.31665277 | 1.61E-13 | hsph1 |
| ENSDARG00000037099 | 2.53560502 | 0.26666363 | 1.19E-18 | irs2a |
| ENSDARG00000038489 | 2.49300998 | 0.22164013 | 5.47E-26 | b3gnt7 |
| ENSDARG00000075833 | 2.44780036 | 0.23852428 | 1.07E-21 | lyve1a |
| ENSDARG00000041022 | 2.39342709 | 0.24258267 | 4.48E-20 | pdcd4b |
| ENSDARG00000060113 | 2.38844383 | 0.21944098 | 2.11E-24 | znf395a |
| ENSDARG00000098925 | 2.29118385 | 0.30319 | 9.07E-12 | prdm1b |
| ENSDARG00000096518 | 2.23143627 | 0.35066543 | 2.20E-08 | si:ch73-95a24.1 |
| ENSDARG00000069282 | 2.18169268 | 0.3028213 | 1.12E-10 | bbc3 |
| ENSDARG00000099002 | 2.14552934 | 0.31031123 | 7.16E-10 | creb5a |
| ENSDARG00000101135 | 2.13959387 | 0.31773682 | 2.18E-09 | si:dkey-85k7.7 |
| ENSDARG00000027529 | 2.13089445 | 0.24266781 | 6.23E-16 | hmox1a |
| ENSDARG00000024746 | 2.1286108 | 0.39057968 | 3.26E-06 | hsp90aa1.2 |
| ENSDARG00000036848 | 2.0836956 | 0.17989377 | 1.55E-27 | slc43a2a |
| ENSDARG00000023217 | 2.02945464 | 0.19637621 | 5.68E-22 | crema |
| ENSDARG00000020761 | 2.01762514 | 0.22426926 | 1.03E-16 | arrdc2 |
| ENSDARG00000094557 | 2.01254617 | 0.32863327 | 9.16E-08 | nupr1 |

B

| ENSEMBL Gene ID | log2FoldChar | lfcSE | padj | gene_name |
|---|---|---|---|---|
| ENSDARG00000053204 | -1.9815825 | 0.28403354 | 4.82E-10 | snx22 |
| ENSDARG00000061923 | -1.9630544 | 0.16346249 | 1.47E-29 | amotl2a |
| ENSDARG00000040764 | -1.829106 | 0.19509112 | 3.97E-18 | id1 |
| ENSDARG00000078800 | -1.6770781 | 0.3338926 | 2.50E-05 | slc26a10 |
| ENSDARG00000037555 | -1.6720874 | 0.3033587 | 2.41E-06 | atoh8 |
| ENSDARG00000042934 | -1.659224 | 0.23592535 | 3.34E-10 | ctgfa |
| ENSDARG00000018441 | -1.5575572 | 0.1724045 | 7.62E-17 | heg1 |
| ENSDARG00000024204 | -1.523809 | 0.22797693 | 2.98E-09 | mcm3 |
| ENSDARG00000045748 | -1.4951781 | 0.18453174 | 1.60E-13 | stab2 |
| ENSDARG00000061948 | -1.4915171 | 0.24854555 | 1.85E-07 | amotl2b |
| ENSDARG00000031894 | -1.4695724 | 0.23481427 | 4.08E-08 | lef1 |
| ENSDARG00000099395 | -1.4649494 | 0.33276355 | 0.000335193 | cables1 |
| ENSDARG00000101707 | -1.4281017 | 0.34863545 | 0.001062347 | si:ch211-156b7.4 |
| ENSDARG00000030104 | -1.4200317 | 0.25455302 | 1.76E-06 | sh3bp4 |
| ENSDARG00000035694 | -1.4188867 | 0.32617363 | 0.00040538 | stm |
| ENSDARG00000038785 | -1.417025 | 0.19775843 | 1.46E-10 | abcf2a |
| ENSDARG00000002216 | -1.3707023 | 0.2790729 | 4.09E-05 | tbx3a |
| ENSDARG00000090337 | -1.3600781 | 0.22591338 | 1.66E-07 | pprc1 |
| ENSDARG00000012506 | -1.3427077 | 0.29865342 | 0.000236107 | rnpep |
| ENSDARG00000038703 | -1.3397517 | 0.31008385 | 0.000453295 | hkdc1 |

C

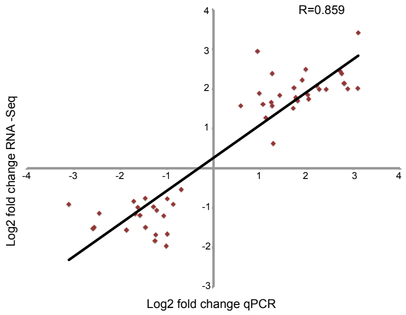


D

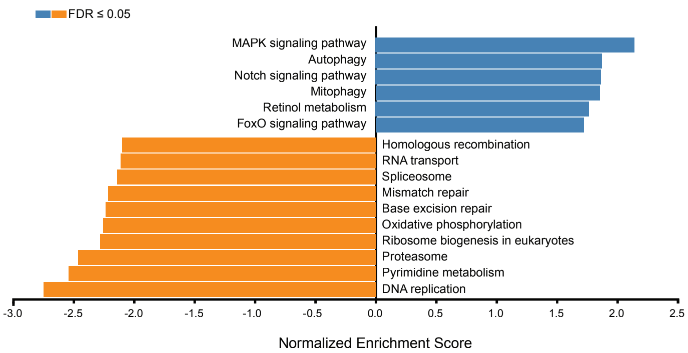


E

| H. sapiens | | D. rerio | | Overlap | |
|---|---|---|---|---|---|
| DE genes | Nr. of genes | DE genes | Nr. of genes | Nr. of genes | % |
| UP | 647 | DOWN | 483 | 30 | 6.2 |
| DOWN | 367 | UP | 584 | 47 | 8.0 |
| ALL | 1014 | ALL | 1067 | 146 | 13.7 |

Tsaryk et al., Supplementary Fig. 6

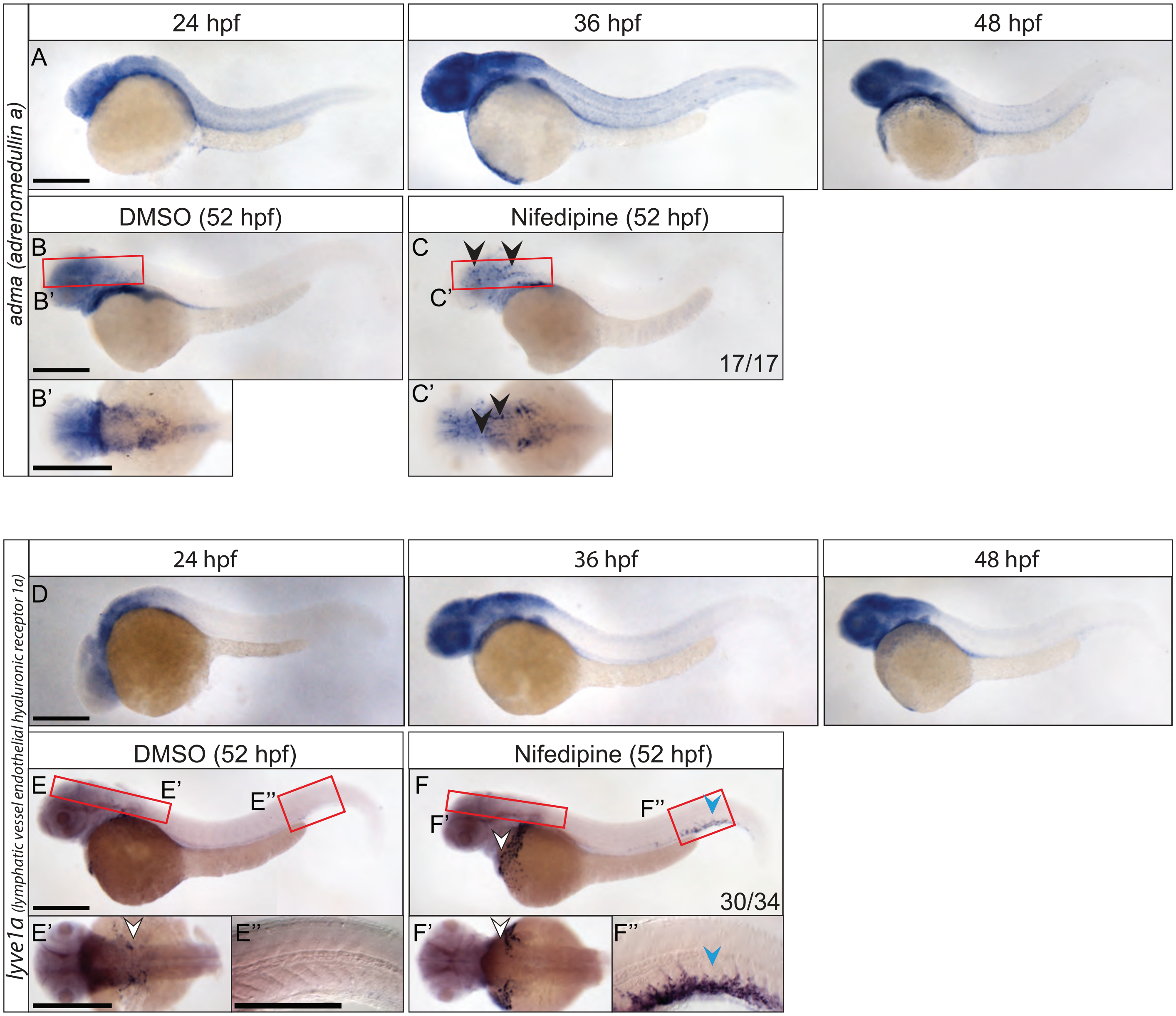


Tsaryk et al., Supplementary Fig. 7

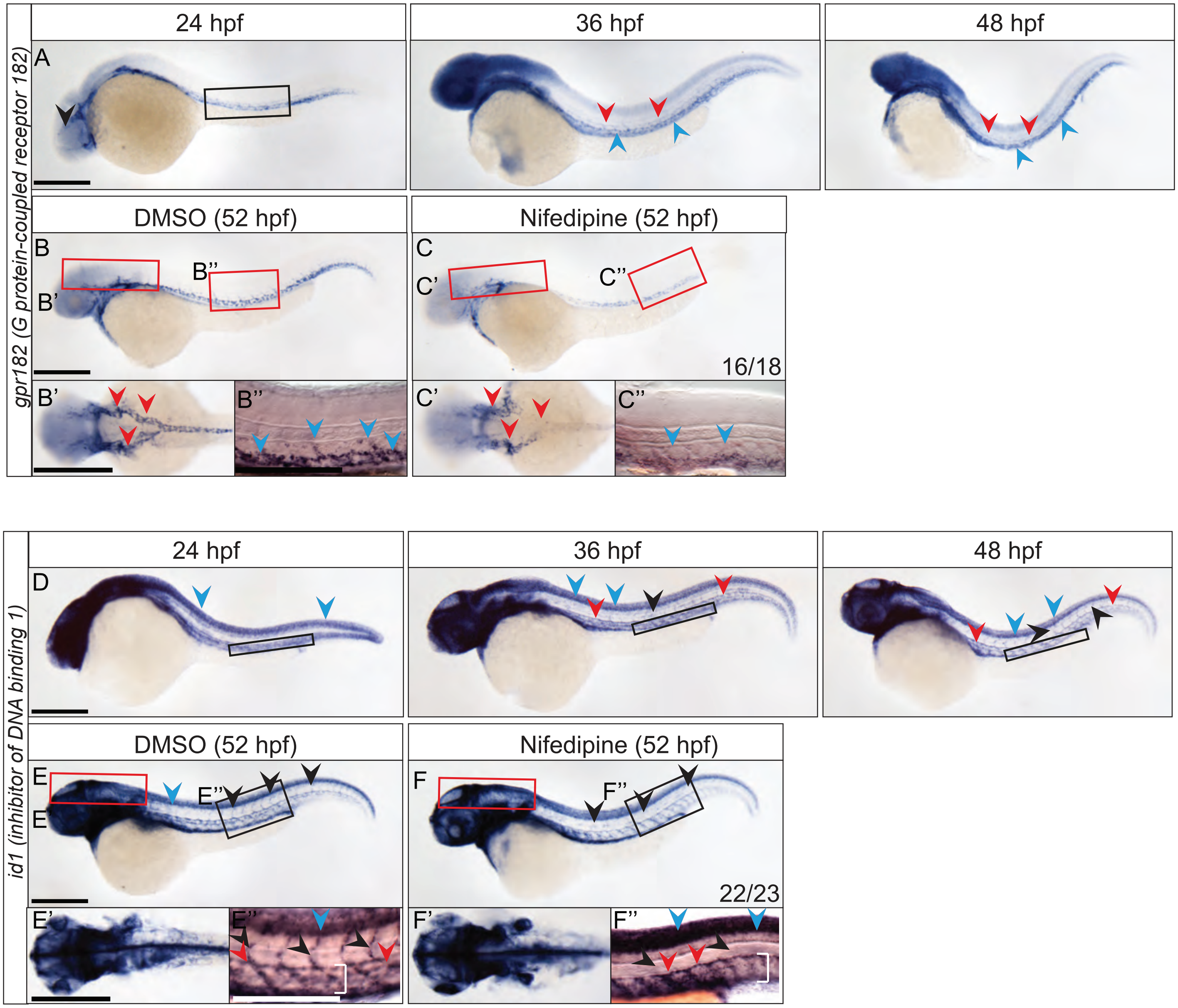


Tsaryk et al., Supplementary Fig. 8

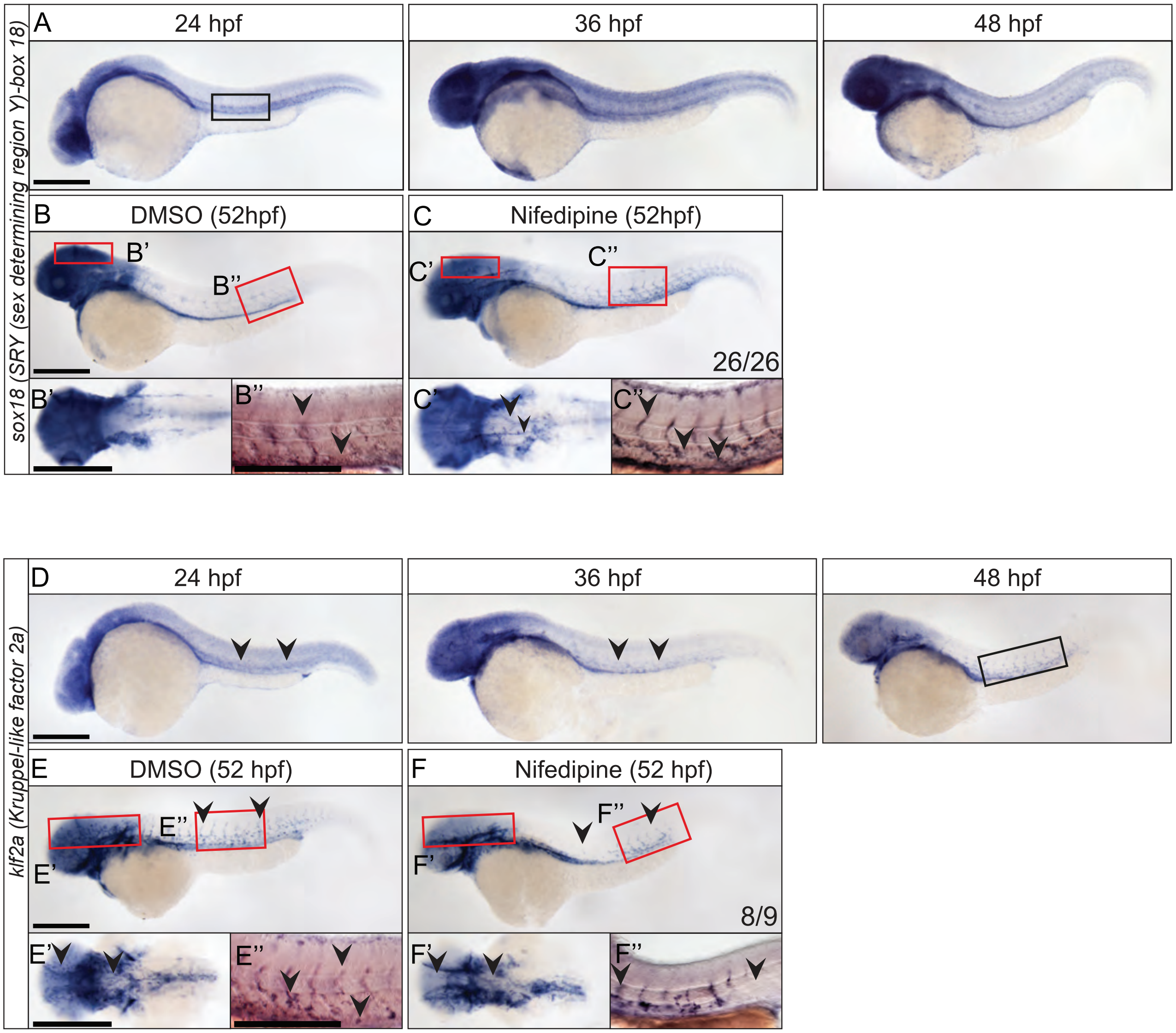


Tsaryk et al., Supplementary Fig. 9
